# Early sex-chromosome evolution in the diploid dioecious plant *Mercurialis annua*

**DOI:** 10.1101/106120

**Authors:** Paris Veltsos, Kate E. Ridout, Melissa A. Toups, Santiago C. González-Martínez, Aline Muyle, Olivier Emery, Pasi Rastas, Vojtech Hudzieczek, Roman Hobza, Boris Vyskot, Gabriel A.B. Marais, Dmitry A. Filatov, John R. Pannell

## Abstract

Suppressed recombination around a sex-determining locus allows divergence between homologous sex chromosomes and the functionality of their genes. Here, we reveal patterns of the earliest stages of sex-chromosome evolution in the diploid dioecious herb *Mercurialis annua* on the basis of cytological analysis, *de novo* genome assembly and annotation, genetic mapping, exome resequencing of natural populations, and transcriptome analysis. Both genetic mapping and exome resequencing of individuals across the species range independently identified the largest linkage group, LG1, as the sex chromosome. Although the sex chromosomes of *M. annua* are karyotypically homomorphic, we estimate that about a third of the Y chromosome has ceased recombining, a region containing 568 transcripts and spanning 22.3 cM in the corresponding female map. Patterns of gene expression hint at the possible role of sexually antagonistic selection in having favored suppressed recombination. In total, the genome assembly contained 34,105 expressed genes, of which 10,076 were assigned to linkage groups. There was limited evidence of Y-chromosome degeneration in terms of gene loss and pseudogenization, but sequence divergence between the X and Y copies of many sex-linked genes was higher than between *M. annua* and its dioecious sister species *M. huetii* with which it shares a sex-determining region. The Mendelian inheritance of sex in interspecific crosses, combined with the other observed pattern, suggest that the *M. annua* Y chromosome has at least two evolutionary strata: a small old stratum shared with *M. huetii*, and a more recent larger stratum that is probably unique to *M. annua* and that stopped recombining about one million years ago.

**Article summary:** Plants that evolved separate sexes (dioecy) recently are ideal models for studying the early stages of sex-chromosome evolution. Here, we use karyological, whole genome and transcriptome data to characterize the homomorphic sex chromosomes of the annual dioecious plant *Mercurialis annua*. Our analysis reveals many typical hallmarks of dioecy and sex-chromosome evolution, including sex-biased gene expression and high X/Y sequence divergence, yet few premature stop codons in Y-linked genes and very little outright gene loss, despite 1/3 of the sex chromosome having ceased recombination in males. Our results confirm that the *M. annua* species complex is a fertile system for probing early stages in the evolution of sex chromosomes.

## Introduction

Separate sexes, or dioecy, have evolved repeatedly from hermaphroditism in flowering plants, with about half of all angiosperm families having dioecious members (Renner and Ricklefs, 1995; Renner, 2014). The evolution of dioecy sets the stage for the possible evolution of sex chromosomes (Charlesworth and Charlesworth, 1978), which originate as a pair of homologous autosomes that may gradually diverge in sequence when recombination between them is suppressed. Such divergence is extreme in many animal species with old sex chromosomes, for which it can be difficult even to identify homologous sequences. In some plants, like *Silene latifolia* (Ono, 1939; Krasovec *et al.*, 2018) and *Rumex hastatulus* (Smith, 1955; Hough *et al.*, 2014), the homologous chromosomes have diverged sufficiently to become heteromorphic and distinguishable by the karyotype. In other plants, like *Asparagus officinalis* (Loeptien, 1979; Telgmann-Rauber *et al.*, 2007) and *Carica papaya* (Horovitz and Jiménez, 1967; Liu *et al.*, 2004), the sex chromosomes remain indistinguishable by the karyotype, and compromised gene function may be mild. Early sex-chromosome evolution usually involves the accumulation of repetitive sequences in a non-recombining region, but also loss of genes, which is presumably preceded by pseudogenization and loss of function.

Dioecious plants provide good models for studying the evolution of sex chromosomes because separate sexes have often evolved recently, potentially allowing us to observe the very earliest stages in sex-chromosome divergence (reviewed in Charlesworth (2016)). To date, the most comprehensive and detailed work on the evolution of plant sex chromosomes comes from the study of species with highly divergent heteromorphic sex chromosomes, such as *S. latifolia* (Kejnovsky and Vyskot, 2010) and *Rumex* species (Navajas-Pérez *et al.*, 2005), i.e., those that have already proceeded down a path of Y-chromosome degeneration and divergence from their X homologue. In these species, it has been possible to measure rates of loss of genes or their function (Krasovec *et al.*, 2018), differences in codon use between homologues (Qiu *et al.*, 2011), and differences in patterns of gene expression at sex-linked loci (Zemp *et al.*, 2016). Such measurements have generally confirmed the idea that non-recombining parts of sex chromosomes quickly degenerate in function, as expected by theory (Lynch *et al.*, 2016). For instance, degeneration may occur through structural genetic changes (Charlesworth *et al.*, 2005), or the accumulation of repetitive elements (Wang *et al.*, 2012b) and deleterious mutations (Charlesworth *et al.*, 2005; Charlesworth, 2013), driven by processes such as background selection (Charlesworth *et al.*, 1993b), selective sweeps (Maynard Smith and Haigh, 1974) or Muller’s Ratchet (Charlesworth *et al.*, 1993a).

Two hypotheses have been proposed to explain the suppression of recombination on the sex chromosomes of plants that have evolved dioecy from hermaphroditism. One hypothesis postulates that dioecy initially evolves through the spread of male-and female-sterility mutations, and that these mutations must become linked on opposite chromosomes to avoid the expression of either hermaphroditism, or both male and female sterility simultaneously (Charlesworth and Charlesworth, 1978). The main experimental support for this two-locus model comes from classic genetic studies in *Silene latifolia* (Westergaard, 1958) that demonstrated the presence of two sex-determining factors on the Y-chromosome, the stamen-promoting factor (SPF) and gynoecium suppression factor (GSF). More recent work mapped the location of SPF and GSF genes on the *S. latifolia* Y-chromosome (Kazama *et al.*, 2016), though the actual GSF and SPF genes have yet to be identified. Nevertheless, despite the likely importance of two loci in the evolution of dioecy, the two-locus model does not readily explain why non-recombining regions on sex chromosomes often expand greatly (Bergero and Charlesworth, 2009), probably well beyond the region harboring the original sex-determining genes.

The second hypothesis invokes expansion of the non-recombining region and divergence of the sex chromosomes with the accumulation of genes that have different effects on the fitness of males and females (Gibson *et al.*, 2002). This generates selection for reduced recombination between the sex-determining allele and nearby alleles with sex-specific effects (Charlesworth, 1991; Charlesworth *et al.*, 2005; Bergero and Charlesworth, 2009), so that alleles with sex-specific fitness benefits (i.e., sexually antagonistic alleles) occur on the sex chromosome that spends most of its time in individuals of the sex to which the benefits apply (Rice, 1987). Mutations that differentially affect gene expression between males and females are also expected to be enriched in the non-recombining region of the sex chromosomes compared with other regions of the genome, as found in *S. latifolia* (Zemp *et al.*, 2016). The suppression of recombination is expected to extend consecutively to generate linkage between the sex-determining locus and more sexually antagonistic loci (Charlesworth, 2015), and these extensions can be identified as discrete ‘strata’, with greater genetic divergence between the sex chromosomes in strata that ceased recombining earliest. Evidence for such strata has been found both in animals (Lahn and Page, 1999; Nam and Ellegren, 2008) and plants (Bergero *et al.*, 2007; Wang *et al.*, 2012b).

If we are to make further progress in understanding the initial suppression of recombination associated with Y-chromosome degeneration, we need to consider systems in which the very earliest stages of sex-chromosome evolution might be observed, either in the youngest evolutionary strata of old (heteromorphic) sex chromosomes, or in sequences close to the sex-determining locus of young homomorphic sex chromosomes that have only just started to diverge. Some progress has been made in studies of several species with homomorphic sex chromosomes, e.g., *Asparagus officinalis* (Loeptien, 1979; Telgmann-Rauber *et al.*, 2007), *Spinacia oleracea* (Yamamoto *et al.*, 2014), *Diospyros lotus* (Akagi *et al.*, 2014), *Fragaria chiloensis* (Tennessen *et al.*, 2016), *Populus* (Geraldes *et al.*, 2015), *Carica papaya* (Horovitz and Jiménez, 1967; Liu *et al.*, 2004) and *Salix* (Pucholt *et al.*, 2015). However, our understanding of the early stages of sex-chromosome evolution remains poor.

Here, we report evidence for the very early evolution of sex chromosomes in the sexually dimorphic dioecious herb *Mercurialis annua* (Euphorbiacae), a plant in which separate sexes have been much studied in the past and that has also become a model to understand evolutionary transitions between combined and separate sexes (Obbard *et al.*, 2006), and the evolution of sexual dimorphism (Sánchez-Vilas *et al.*, 2010). Until recently, gender in *M. annua* was thought to be determined by allelic variation at three independent loci (Durand *et al.*, 1987; Durand and Durand, 1991), but it is now known to have a simple XY system (Khadka *et al.*, 2005; Russell and Pannell, 2015). The sex chromosomes in *M. annua* are homomorphic (Durand, 1963), and crosses between males with ‘leaky’ gender expression reveal that YY males are viable but partially sterile (Kuhn, 1939). It thus seems that the *M. annua* Y chromosome is in the early stages of degeneration.

Our study is based substantially on *de novo* assembly and annotation of the diploid *M. annua* genome (2n = 2x = 16). Whole-genome data for species with separate sexes are scarce in plants, with the few exceptions being *Carica papaya* (Liu *et al.*, 2004), *Vitis* (Fechter *et al.*, 2012), *Diospyros lotus* (Akagi *et al.*, 2014), and *Populus* (Tuskan *et al.*, 2006), which have largely homomorphic sex chromosomes (Filatov, 2015). For the genome assembly of *M. annua*, we combined short-read sequencing on the Illumina platform with long-read technology developed by Pacific Biosciences. We also analyzed the karyotype of *M. annua* using current imaging technology in an attempt to identify heteromorphism at the cytological level, and we obtained SNPs from the transcriptomes of small families to construct a genetic map for *M. annua*, to identify non-recombining (sex-linked) genes and scaffolds, and to compare them with non-sex-linked regions. We then investigated the sex chromosomes with regard to evolutionary strata and degeneration, including the fixation pseudogenising mutations in the inferred fully Y-linked sequences, and the proportion of genes deleted from the Y. Finally, we examined sex-biased gene expression and assessed whether sex-biased genes might be enriched on the sex chromosomes, as expected by theory (Connallon and Clark, 2010; Meisel *et al.*, 2012). Our results suggest that the sex chromosomes of *M. annua* are about 1.5 million years old. Early stages of Y-chromosome degeneration are clearly apparent, but there are also signs that patterns of gene expression might have been affected by the accumulation of sexually antagonistic mutations.

## Materials and Methods

### Hairy root culture, chromosome preparation and cytogenetics

*Mercurialis annua* seeds were sterilized by incubation in 4% sodium hypochloride and subsequently washed in 50% ethanol and sterile water. Seeds were grown on MS medium and leaf discs were taken from 2-3 week old plants, which had been sexed using a previously developed Sequence Characterised Amplified Region (SCAR) marker (Khadka *et al.*, 2002). *Agrobacterium rhizogenes* strain ARqua1 (Quantdt *et al.*, 1993) was grown overnight at 28°C in LB medium supplemented with 300 µM acetosyringone to OD_600 = 0.6. The culture was resuspended in liquid 1/2 MS with 300 µM acetosyringone and used for direct inoculation of *Mercurialis annua* leaf discs. Explants were cocultivated at 28°C on MS medium with 300µM acetosyringone for 2 days. After cocultivation, explants were moved to MS medium supplemented with 300µg/L Timentin. Media were changed every 2 weeks.

In order to synchronize the hairy roots of *Mercurialis annua*, the DNA polymerase inhibitor aphidicolin was added for 12 h. Mitoses were then accumulated in protoplasts using oryzalin treatment, transferred on chromic acid washed slides by dropping, and stored at −20 ºC until use. For FISH experiments, slides were denatured in 7:3 (v/v) formamide: 2xSSC for 2 min at 72 ºC, immediately dehydrated through 50%, 70%, and 100% ethanol (−20 ºC), and air dried. The hybridization mixture (30 µl per slide) consisted of 200 ng of labeled probe, 15 µl formamide, 6 µl 50% dextrane sulfate, and 3 µl of 20x SSC. The volume was brought to 30 µl by adding TE, pH 8. The probes were denatured at 70ºC for 10 min, and slides hybridized for 18 h at 37ºC in a humid chamber. Slides were analyzed using an Olympus Provis AX70 microscope, and image analysis was performed using ISIS software (Metasystems). DNA was labeled with Fluorolink Cy3-dUTP (Amersham Pharmacia Biotech; red labeling) using a Nick Translation mix (Roche) or with SpectrumGreen direct-labeled dUTP (Vysis; green labeling) and a Nick Translation kit (Vysis).

### Genome and transcriptome sequencing

Diploid *Mercurialis annua* individuals from north-western France were grown together in the glasshouse. DNA for genomic sequencing was taken from leaf tissue of a single male, M1. RNA samples were collected from leaves and flower buds of this individual as well as from three females, G1, G2 and G3, and two additional males, M2 and M3, all of which were unrelated. 65 F_1_ and F_2_ progeny (five families) were then produced by crossing G1xM1 and G2xM1 (F_1_) as well as three pairs of G1xM1 progeny (F_2_) (Table S1), which were also used for RNA extraction and transcriptome sequencing. DNA was extracted using a Qiagen Plant DNeasy kit (Qiagen, Hilden, Germany). Illumina paired-end and mate-pair sequencing was carried out by the Beijing Genomics Institute (BGI) using Illumina HiSeq 2000 technology (100 bp reads). Pacific Biosciences long-read sequencing was performed on the individual M1 by the Centre for Integrative Genomics hosted at the University of Lausanne. RNA was extracted from a mixture of flower buds and leaf tissues using the Qiagen plant RNAeasy kit (Qiagen, Hilden, Germany), and individual libraries were prepared for all 65 individuals (Table S1), which were sequenced on three lanes of Illumina HiSeq 2000 at the Wellcome Trust Centre for Human Genetics, Oxford.

### Transcriptome assembly and annotation

The SEX-DETector pipeline (http://lbbe.univ-lyon1.fr/-SEX-DETector-.html?lang=fr) was used to produce the reference transcriptome (henceforth ‘ORFs’), and obtain allele-specific expression and SNPs for X and Y haplotypes. Sex linkage was inferred from genetic mapping which used more data (see below). To identify sex-linked haplotypes, we employed a probabilistic model based on maximum-likelihood inference, implemented in SEX-DETector (Muyle *et al.*, 2016). SEX-DETector is based on the idea that X-linked and Y-linked haplotypes are passed only from fathers to daughters or from fathers to sons, respectively. SEX-DETector is embedded into a Galaxy workflow pipeline that includes extra-assembly, mapping and genotyping steps prior to sex-linkage inference, and has been shown to have greater sensitivity, without an increased false positive rate, than without these steps (Muyle *et al.*, 2016). First, poly-A tails were removed from transcripts using PRINSEQ (Schmieder and Edwards, 2011), with parameters -trim_tail_left 5 -trim_tail_right 5. rRNA-like sequences were removed using riboPicker version 0.4.3 (Schmieder and Edwards, 2011) with parameters -i 90 -c 50 −l 50 and the following databases: SILVA Large subunit reference database, SILVA Small subunit reference database, the GreenGenes database and the Rfam database. Transcripts were then further assembled within Trinity components using cap3 (Huang and Madan, 1999), with parameter -p 90 and custom Perl scripts. Coding sequences were predicted using Trinity TransDecoder (Haas *et al.*, 2013), including PFAM domain searches as ORF retention criteria; these sequences were taken as our reference transcriptome. The RNAseq reads from parents and progeny were mapped onto the reference transcriptome using BWA (Li and Durbin, 2009). The alignments were analyzed using reads2snp, a genotyper for RNAseq data that gives better results than standard genotypers when X and Y transcripts have different expression levels (Tsagkogeorga *et al.*, 2012).

The SEX-DETector transcripts were mapped onto the genome using GMAP v2017-06-20 (Wu and Watanabe, 2005), disallowing chimeric genes, mapping 38,963 (99%) of them to 48,298 loci. We then collapsed the inferred ORFs into 34,105 gene models using gffread v0.9.9 (http://ccb.jhu.edu/software/stringtie/gff.shtml). In total, 26,702 (67.9%) of the ORFs could be annotated using fast blastx on the Swiss-Prot database and Blast2GO (Conesa *et al.*, 2005) in Geneious v9 (Kearse *et al.*, 2012). The resulting transcriptome annotation information, including GO terms, is presented in File S4. We characterized genes as candidate sex-determining genes by identifying those containing the following strings in their annotation: flower, auxin, cytokinin, pollen, kelch, ethylene, swi, retinol, calmodulin, short-root, jasmonic. This manual characterization allowed us to visualize the mapping locations, test for their enrichment in the non-recombining region, and identify whether any were among the outliers in various transcript-associated metrics.

### Genome size estimation, assembly and annotation

The genome size was estimated from the distribution of k-mers (size 31) obtained from jellyfish v2.1.0 (Marçais and Kingsford, 2011) and analyzed by the web version of GenomeScope v1 (Vurture *et al.*, 2017), limiting max k-mer coverage to 1000.

Sliding-window trimming and adaptor removal were carried out using Trimmomatic v. 0.30, with the default parameters (Bolger *et al.*, 2014). Exact duplicate read pairs were collapsed using fastx-collapser from the Fastx-Toolkit (http://hannonlab.cshl.edu/fastx_toolkit/). Low complexity masking was carried out using DUST (Morgulis *et al.*, 2006), with the default parameters, and ambiguous reads were removed. Pacific Biosciences long reads were error-corrected using Bowtie2 version 2.1.0 (Langmead and Salzberg, 2012) in combination with LSC version 0.3.1 (Au *et al.*, 2012).

Filtered paired-end and mate-paired genomic reads were assembled using SOAPdenovo2 (Luo *et al.*, 2012), with k-mer values between 35 and 55 (odd values only). The best assembly was chosen using REAPR (Hunt *et al.*, 2013). GapCloser (Luo *et al.*, 2012) was run on the best assembly to correct false joins and fill gaps. Error-corrected Pacific Biosciences reads were then used to extend scaffolds, to fill gaps, and to join scaffolds using PBJelly2 (English *et al.*, 2012). Additional scaffolding was carried out using default parameters on SSPACE (Boetzer *et al.*, 2011), which revisits gaps using existing paired-end and mate-paired sequences. This step is intended to correct for any ambiguity introduced by the low-coverage of PacBio reads. Finally, L_RNA_Scaffolder (Xue *et al.*, 2013) was used to bridge genomic scaffolds using the transcript assembly.

Transposable elements (TEs) and tandem repeats were predicted using a combination of Tandem Repeats Finder (Benson, 1999), RepeatModeler and RepeatMasker (Smit *et al.*, 2013). Repeat libraries from *Mercurialis* (from RepeatModeler), Euphorbiacae and *Vitis vinifera* were used for masking the genome before further analyses.

### Summary of SNP calling pipelines

We called SNPs using four different pipelines. SNPs were called on RNAseq data from the two largest families by SEX-DETector, to call X-and Y-specific SNPs, and infer haplotypes with premature stop codons and allele-specific expression. SNPs were also called from genome capture data of multiple diploid populations, for independent analysis of genomic regions of differing sex linkage as inferred from the genetic map. In addition, SNPs were called on all individuals from five families, to perform genetic mapping. Finally, SNPs were called on transcripts from the three unrelated males (M1, M2 and M3) and the three unrelated females (G1, G2 and G3) for comparisons between ORFs of differing sex linkage and comparisons of pairwise diversity between species. The subset of these for which there was evidence of separate X and Y copies, was analyzed in a phylogenetic framework to estimate the (tree-based) evolution rate of the Y sequences. Specific details of each pipeline are given in the corresponding sections below.

### Genetic map construction

We called SNPs independently of the SEX-DETector pipeline, using all individuals available over the three generations of our family crosses. We first mapped RNAseq reads to the reference transcriptome using Bowtie2 v2.3.1 (Langmead and Salzberg, 2012). The resulting sam files were converted to bam, sorted and converted to mpileup format using samtools v 1.3 (Li *et al.*, 2009). The resulting posterior file containing segregation information of transcripts in the five mapping families (>164,000 markers and their genotype likelihoods) was taken through the LepMap3 (LM3) pipeline (Rastas, 2017). First, we calculated the relatedness between individuals using the IBD module (of LM3) using a random subset of 3,000 markers. Three individuals were discarded because their relatedness to their putative father (M6) was < 0.2, i.e., they were likely the result of a different cross through contamination. The parental genotypes were then called using ParentCall2, with parameter halfSibs=1, to take into account the half-sib family structure, in addition to the genotypic information of offspring, parents and grandparents. The 158,000 remaining markers were further filtered using the Filtering2 module, with parameter dataTolerance=0.001, to remove markers segregating in a distorted fashion. We then combined markers of each transcript into pseudo-markers, as they were not informative enough on their own to separate into linkage groups. This was done by running OrderMarkers2 separately for each transcript without recombination (parameter values: recombination1=0, recombination2=0), and then obtaining genotype likelihoods for each transcript with parameter outputPhasedData=4. This resulted in 13,261 markers, each consisting of a single transcript, which we re-formatted as input for LM3. We defined linkage groups by running SeparateChromosomes2 with default parameters (lodLimit=10), adding additional markers to the resulting 8 linkage groups using JoinSingles2 with lodLimit=8 and lodDifference=2. We were able to assign 10,076 transcripts into eight linkage groups, which were ordered using OrderMarkers2. The linkage maps were drawn with R/qtl v1.41 (Broman *et al.*, 2003) running in R 3.4.2 (R Development Core Team, 2007).

### Sequence divergence in *M. annua*, and in comparison with closely related species

We aligned reads from six unrelated individuals from France (M1, M2, M3, G1, G2, G3) to the 39,302 open reading frames from the reference transcriptome assembled via the SEX-DETector pipeline, based on the M1 male transcriptome and using the BWA-MEM algorithm of BWA v0.7.13 (Li and Durbin, 2009). Picard Tools v2.2.1 (http://broadinstitute.github.io/picard/) were used to mark duplicate read pairs. Local realignment around insertions and deletions (indels) was performed with GATK v3.7 (DePristo *et al.*, 2011), followed by SNP calling on each individual using the HaplotypeCaller module in GATK (McKenna *et al.*, 2010). Joint SNP calling was performed using GATK’s GenotypeGVCFs module, overlapping SNPs and indels were filtered out with the VariantFiltration module, and SNPs and indels were separated and filtered to produce two high-quality variant sets with the following parameters in the VariantFiltration module: ‘QUAL < 30’ ‘DP < 30’ ‘MQ0 >= 4 && ((MQ0 / (1.0 * DP)) > 0.1)’ ‘QD < 5.0’. The high-quality SNP set was used to perform variant quality score recalibration to filter the full SNP set. This set was used to calculate within-species nucleotide diversity (π) using vcftools v 0.1.13 (Danecek *et al.*, 2011).

Pairwise d_N_ and d_S_ values from *M. huetii* and *R*. *communis* were obtained using yn00 from PAML 4.9 (Yang, 2007). This analysis used 4,761 and 2,993 homologues from the *M. huetii* and *R. communis* (http://castorbean.jcvi.org/introduction.shtml) transcriptomes, based on a *de novo* assembly using default settings in Trinity (Haas *et al.*, 2013; File S5). Sequences were aligned with LASTZ (Harris, 2007). Effective codon usage was calculated using ENCprime (https://jnpopgen.org/software/).

To estimate the rate of evolution of Y sequences, we identified one-to-one orthologs between the *M. annua* sex linked genes and the closely related species *M. huetii* and *R. communis*, through reciprocal best BLASTP, with an e-value of 1e^-50^, a culling limit of 1, and considering only transcripts that were not split across contigs. The 98 orthologs identified were aligned with LASTZ (Harris, 2007) and analysed with PAML 4.9 (Yang, 2007) to obtain tree-based estimates of d_N_ and d_S_ values. We employed three models: the ‘null model’ (M_0_), where all parts of the tree are assumed to have the same d_N_/d_S_values; the ‘branch-specific’ model (M_1_), where each branch may have its own d_N_/d_S_value; and the ‘two-ratio’ model (M_2_), where the branch of interest (Y) may have a d_N_/d_S_ ratio that differs from the rest of the tree (Nielsen and Yang, 1998). Preliminary analysis revealed similar results for the M_1_ and M_2_ models (results not shown); we thus chose the M_2_ as the simpler model for further analysis. We tested whether the Y branch evolved differently from the rest of the tree, using a likelihood ratio test (LRT) between the M_2_ and M_0_ models, with one degree of freedom and LRT = 2 x abs(l_2_-l_0_), where l_2_ and l_0_ are the likelihoods for M_2_ and M_0_respectively.

### Expression analysis

Reads from the RNA libraries of all 30 females and 35 males were pseudoaligned to the reference transcriptome using Kallisto v0.43.1 (Bray *et al.*, 2016). The resulting raw count data were compiled in a table, and differential expression analysis between male and female samples was conducted with edgeR v3.18.1 (Robinson *et al.*, 2010), running in R v.3.4.0 (R Development Core Team, 2007). Genes were filtered from the analysis if their average log_2_ count per million (as computed by edgeR’s aveLogCPM function) was negative. This retained genes with an average count of 24 or more per sample. Two individual libraries were removed that were obvious outliers in the MDS plots of the filtered data. Libraries were normalized with the default (TMM) normalization. Dispersion was measured with default parameters using a negative binomial model. Sex-biased transcripts were defined statistically, i.e. only by a false discovery rate (FDR – Benjamini and Hochberg; 1995) of 5%. A statistical definition for sex-biased genes is justified because our RNAseq analysis was based on atypically many samples, and because the sex-biased categories (male-, female-and unbiased) used in our analysis capture biological information in that they differed statistically in various measurements. This would also ensure that our results for effects of sex bias should be robust, because our approach would dilute “real” sex-biased genes with unbiased genes. We note, however, that the lack of minimum log_2_FC in our definition of sex-bias means that we cannot differentiate between allometric differences in tissue composition or gene regulation differences between males and females (Montgomery and Mank, 2016).

SEX-DETector calculates X-and Y-allele expression from each male. X and Y read numbers were summed for each contig and individual separately and divided by the number of X/Y SNPs of the contig and adjusted for the library size of the respective individual. X and Y expression levels were then averaged among individuals, and the ratio of the means was computed. Gene ontology (GO) enrichment analysis was performed on the sex-biased genes, as well as separately for the male-and female-biased genes, using topGO with the weight01 algorithm, which accounts for GO topology (Alexa and Rahnenfuhrer, 2010).

### Comparison of expression between sex-linked genes and autosomal genes

We categorized transcripts based on their genomic location into fully sex-linked (SL; transcripts in the non-recombining region of LG1 in males), pseudo-autosomal (PAR; transcripts in the rest of LG1 that showed recombination in both sexes) and autosomal (Au; transcripts that mapped to the remaining seven linkage groups). We investigated the effects of sex linkage, sex-biased expression and their interactions through linear models using R v 3.4.2 on the following metrics: Absolute male/female log_2_FC (fold-change, i.e., overall sex-bias intensity); nucleotide diversity (π); d_N_/d_S_from pairwise comparisons with *M. huetii* and *R. communis*; and Y/X allele-specific log_2_FC obtained from SEX-DETector. The data were not normally distributed and were thus Box-Cox-transformed after adding 1e^-11^ to all values to allow inclusion of zero values and estimating the transformation parameter using the command boxcoxnc with the Anderson-Darling method in the R package AID v2.3 (Dag *et al.*, 2014). Variables Absolute male/female (log_2_FC) and π remained non-normal after transformation when unbiased genes were included in the analysis, because there were too many transcripts with ~ 0 values. Exclusion of these transcripts’ values did not qualitatively affect the results (data not shown). We also analyzed the sex-biased genes on their own within the same statistical framework. This allowed for both a more similar distribution between the sex-bias categories, and for successful tests of normality after Box-Cox transformation. The summary tables from the models were generated using the R package sjPlot v 2.4 (Lüdecke, 2017).

We also compared the proportions of sex-biased/unbiased genes and male-biased/female-biased genes between pairs of genomic regions of differing sex linkage using chisq tests (or Fishers’s exact tests when the gene numbers were small), employing the sjt.xtab function of the in R package sjPlot v 2.4 (Lüdecke, 2017). The pairs of genomic regions compared were: PAR-SL; Au-LG1; and Au-SL. Finally, we looked for enrichment of candidate genes in the same genomic region comparisons using the same methodology.

### Divergence between males and females from natural populations

To assess divergence between males and females, we examined *F*_ST_ and sex-biased SNP density from individuals sampled from across the species’ range. Seeds obtained from multiple *M. annua* populations (Table S2) were germinated and grown in a glasshouse at the University of Lausanne. Twenty individuals of each sex were selected for genomic analysis to achieve a balance between representation of many populations and sufficient common sequences of good quality. High-quality genomic DNA was extracted from about 100 mg of frozen leaf material using the DNeasy Plant Mini Kit (Qiagen, Hilden, Germany), following standard protocols. Based on the successful probes from a previous gene-capture experiment (see González-Martínez *et al.*, 2017), about 21 Mbp of the *M. annua* diploid genome (175,000 120-mer probes) were sequenced, with sequence capture using SureSelect DNA Capture technology (Agilent Technologies, (Santa Clara, CA) followed by sequencing using Illumina HiSeq 2500, outsourced to Rapid Genomics (Gainesville, FL). Our gene-capture experiment included both genic and intergenic regions and avoided repetitive regions and organelle genomes. Raw data was provided by Rapid Genomics as FASTQ files.

Sequencing quality was assessed using FASTQC v.10.5 (www.bioinformatics.babraham.ac.uk/projects/fastqc). Adapter removal and trimming was done using Trimmomatic v0.36 (Bolger *et al.*, 2014), with options ILLUMINACLIP: TruSeq3-PE.fa:2:30:10 LEADING:3 TRAILING:3 SLIDINGWINDOW: 4:15 MINLEN:36. After filtering, we retained an average of 13 million paired-end sequences per sample. We aligned sequences to the *M. annua* scaffolds using BWA-MEM v.0.7.12 (Li and Durbin, 2009) and processed the alignments with Picard Tools v.1.141 (http://broadinstitute.github.io/picard/). We ran Samtools mpileup (Li *et al.*, 2009) with the probabilistic alignment disabled, and called SNPs using Varscan (Koboldt *et al.*, 2009) with a minimum variant allele frequency of 0.15 and a minimum threshold for homozygotes of 0.85. We required a minimum of 10 reads per site and a Phred quality score > 20.

The gene capture ORFs were aligned to the genomic scaffolds using BLAST v2.2.31 (Altschul *et al.*, 1990). We retained hits with a maximum e-value of e^-20^, and a minimum identity of 95%. In the case of multiple ORFs mapping to the same genomic location, we selected the longest alignment, and removed all overlapping ORFs. This resulted in 21,203 transcripts aligning to the genome, 8,317 of which were also included in the linkage map. All scaffolds with a transcript were assigned a linkage group position in the map. 31 scaffolds contained transcripts assigned to more than one linkage group; in these cases, we assigned the scaffold to the linkage group with >50% of the transcripts (in the case of equal assignment, we excluded the scaffold from the analysis). In total, we were able to assign 6,458 scaffolds to linkage-group positions.

We used the BLAST output to construct a bed file that defined the coordinates of each transcript on a genomic scaffold. We then extracted genetic information for the transcripts specified in the bed file. Using this information, we calculated two complementary measures of differentiation between males and females for transcripts exceeding 100 bp. First, we computed *F*_ST_for each transcript using VCFtools v.0.1.15 (Danecek *et al.*, 2011). Similarly, we assessed sex-biased heterozygosity for each transcript (Toups *et al.*, 2019), which we defined as the log_10_of the male:female SNP-density ratio. We expect this ratio to be zero for autosomal transcripts, and elevated on young sex chromosomes due to excess heterozygosity in males. For transcripts on older sex chromosomes, the X and Y alleles may have diverged substantially, so that the Y alleles would no longer map to the X allele reference. In this case, there will be inflated female heterozygosity, due to X-chromosome hemizygosity in males.

For both *F*_ST_and sex-biased heterozygosity in the two sexes, we assessed differences between the autosomes, PAR, and sex-linked region using Wilcoxon signed rank tests. We then computed moving averages in sliding windows of 20 transcripts on all chromosomes using the rollmean function, from package zoo v1.8-0 in R v3.2.2. In order to identify regions of elevated *F*_ST_ and sex-biased heterozygosity, we computed 95% confidence intervals on the basis of an average of 30 consecutive transcripts on the autosomes calculated 1,000 times. Finally, for both metrics, we assessed whether the top 5% of transcripts were overrepresented on LG1 using chi-squared tests.

### Data Availability

All raw DNA and RNA sequence data generated in this study have been submitted to NCBI under accession SRP098613 BioProject ID PRJNA369310. Scripts, supplementary files and intermediate files are available at https://osf.io/a9wjb/?view_only=fd59f4dbce5a4c4a88ba31036b51fb35 File S1 contains the genetic map data and other information on each mapped transcript. File S2 contains the Sanger sequence of clones containing the X and Y variants, where there is a premature stop codon on the Y, and the script used to make the phylogenetic tree. File S3 summarizes the GO results for sex-biased genes. File S4 contains transcriptome annotation information. File S5 contains the assembled transcriptome and X and Y haplotype predictions from SEX-DETector. File S6 contains the genome and predicted genes on it. File S7 is the *M. annua* repeat library.

## Results

### Male karyotype

A karyotype for a male individual is shown in Figure 1, and chromosome spreads from both sexes are shown in Figure S1. Idiogram data, chromosomal arm ratios and relative length measurements from 50 images (obtained from 5 individuals; Table S3) suggest that all *M. annua* chromosomes can be discriminated from one another. A partial distinction between the one large, two medium and five smaller chromosomes is possible on the basis of the two common ribosomal DNA probes (45S and 5S), with a secondary constriction often visible at the location of the 45S locus. Our analysis was unable to discern any heteromorphic chromosomes, confirming the similar conclusion reached by Durand (1963).

**Figure 1:**
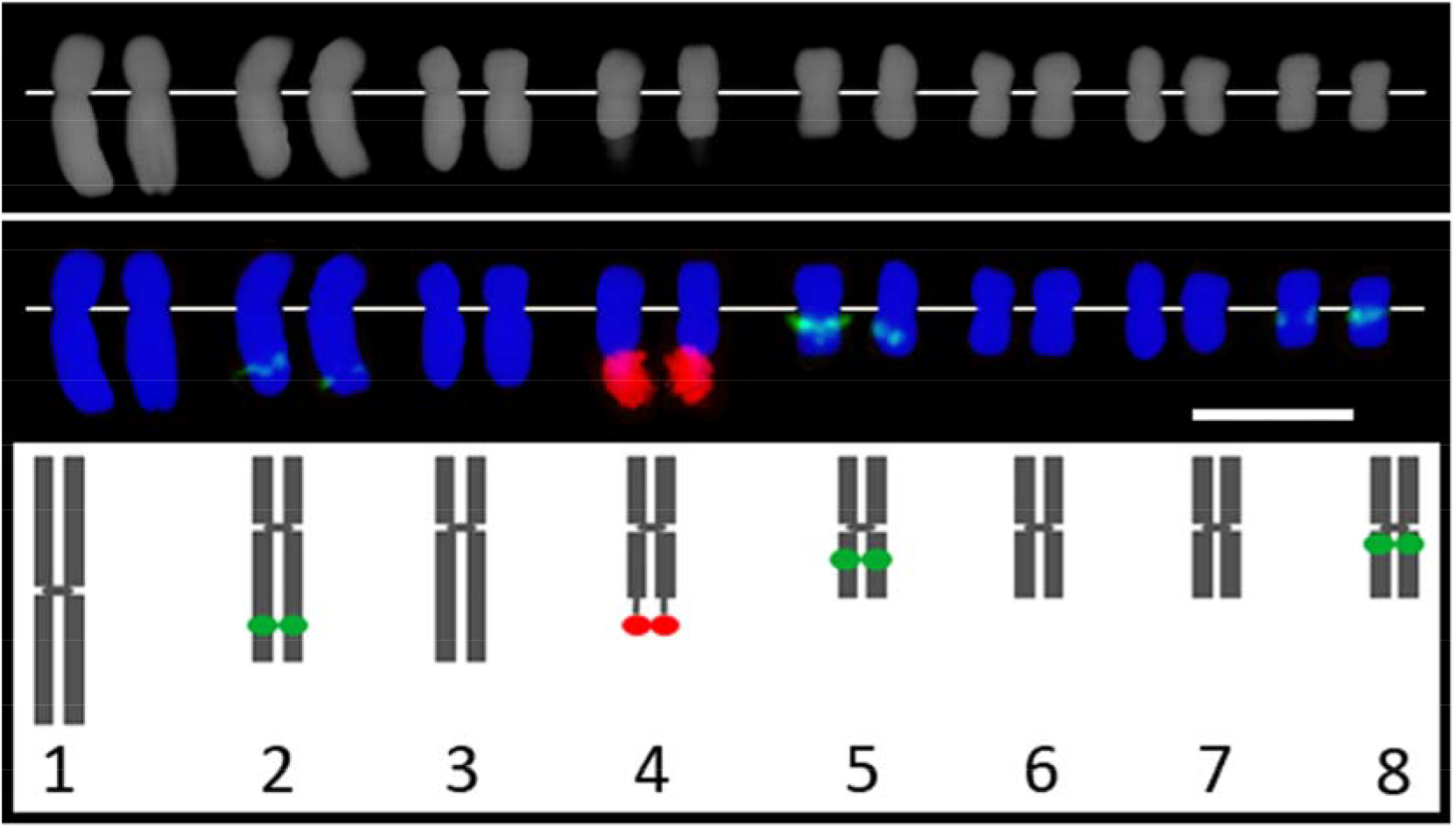
Basic karyotype for a *Mercurialis annua* male individual. The top row shows DAPI counterstain. The middle row shows chromosomes counterstained with DAPI (blue) after bicolor FISH with 45S rDNA (red) and 5S rDNA (green). The bottom row shows an idiogram of the male chromosomes. The white bar in the middle row represents 5 µm.

### *M. annua* genome assembly

Using k-mer distribution, we estimate the haploid genome size to be 325 Mb (Figure S2, Table S4), which is similar to the previous estimate of 640 Mb for the diploid genome (Obbard *et al.*, 2006). Using a combination of short-read Illumina and long-read Pacific Biosciences DNA sequencing, we generated ~56.3 Gb of sequence data from the *M. annua* male M1, corresponding to ~86.6 coverage for a diploid genome size of 650 Mb. After read filtering, genome coverage dropped to ~63.7x (Table S5). *De novo* assembly and scaffolding yielded a final assembly of 89% of the genome (78% without gaps), 65% of which was assembled in scaffolds > 1 kb, with an N_50_ of 12,808 across 74,927 scaffolds (Table S6). Assembly statistics were consistent with those for other species of the Malpighiales (Table S7), as was our estimate of total genomic GC content (34.7%; Smarda *et al.*, 2012). The *M. annua* assembly encompassed over 89% of the assembled transcriptome; most of the unassembled genome sequence data is therefore expected to be repetitive. Using BUSCO2 (Simão *et al.*, 2015), we recovered 76.1% of the 1,440 genes in the BUSCO embryophyte database (of which 3.9% were duplicated). A total of 10.3% of the genes were fragmented, and 13.6% were missing.

Repeat masking identified simple tandem repeats in over 10% of the assembly. This proportion is likely to be an underestimate, given the difficulty in assembling microsatellites. We characterized 15% of the assembly on the basis of homology information and DNA transposon and retrotransposon masking, with an additional 33% corresponding to 1,472 predicted novel transposable elements. The most frequent transposable repeats were *gypsy* LTR, *copia* LTR, and L1 LINE retrotransposons (Table S8), similar to findings for other plant genomes (Chan *et al.*, 2010; Sato *et al.*, 2011; Wang *et al.*, 2012a; Rahman *et al.*, 2013). Across all data, 58% of the ungapped *M. annua* assembly was repetitive (Table S8), corresponding to 44% of the genome. Given that our assembly covered 78% of the genome, and if we assume that the missing fraction comprises only repeats, up to 66% of the *M. annua* genome may be repetitive. This too would be consistent with what is known for other plants, which have a similarly high AT-rich repeat content and a similar number of unclassified repetitive elements (e.g. Chan *et al.*, 2010). The *M. annua* genome appears to be about 240 Mb larger than that of its relative *Ricinus communis* (see Table S7), perhaps reflecting on-going transposon activity.

### Sex-linked transcripts and genetic map for diploid *M. annua*

Genetic mapping recovered eight linkage groups (LG1 through LG8), corresponding to the expected chromosome number for diploid *M. annua* (Durand, 1963). LepMap3 (LM3) identified 568 sex-linked (SL) transcripts, of which 365 were also identified as sex-linked by SEX-DETector and phased into X and Y haplotypes. The 568 SL transcripts mapped to LG1, which is thus the sex chromosome. An additional 1,209 transcripts mapped to the two ends of LG1, and represent two putative pseudoautosomal regions (PAR1 + PAR2; which we refer to these as PAR and consider them together in all analyses); 8,299 transcripts mapped to the other (autosomal) linkage groups. In total, 641 markers could be resolved from one another and were thus informative. We inferred the sex-linked region to comprise of transcripts without recombination in males and in the same phase as that of their paternal grandfather. Assuming an equal transcript density across the genome, the sex-linked region represents 32.6% (about a third) of LG1 and 5.8% of the 320 Mb haploid genome (Obbard *et al.*, 2006), or 18.6 Mb. This Y-linked region maps to LG1 at male position 53.85 cM and spans 22.23 cM on the female recombination map (from positions 46.44 - 68.67 cM; Figure 2). The total male and female recombination maps were 700.43 and 716.06 cM, respectively. All marker names and their associated positions and metrics are provided in File S1.

**Figure 2:**
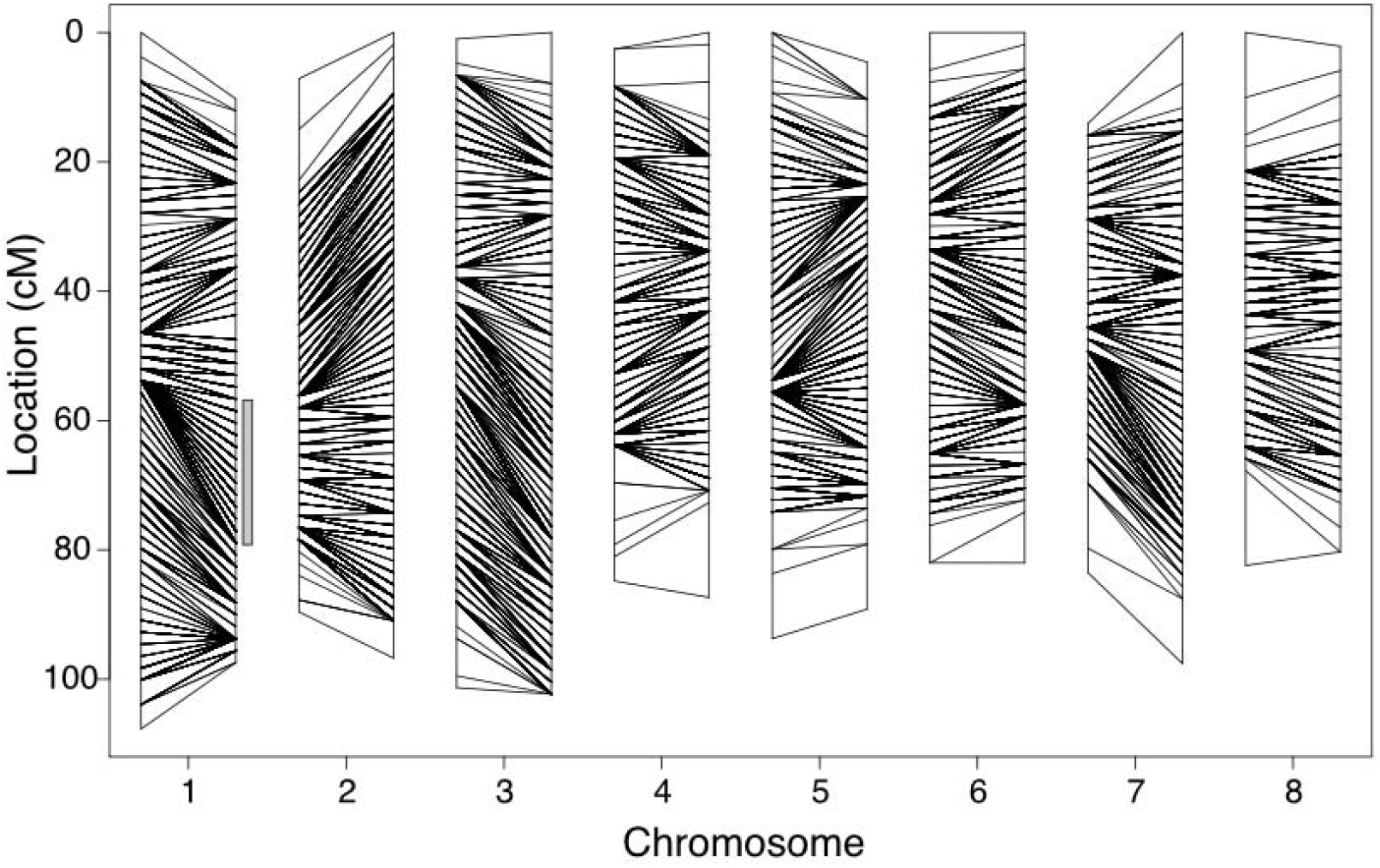
Genetic map of the 8 linkage groups (LG) for *M. annua*, with each LG represented by the male (left) and female (right) maps. Lines linking the two maps indicate the mapping position of groups of transcripts that segregated together (10,076 total transcripts). The sex-linked region is at 53.85 cM of chromosome 1 (LG1) on the male recombination map, and is highlighted by the gray vertical bar. The 2 pseudoautosomal regions (PAR) comprise the rest of LG1.

### Characterization of contigs by sex linkage and ORF localization

We characterized the genome assembly by mapping all ORFs using GMAP and dividing genomic contigs into three groups: ‘sex-linked’, ‘autosomal’, and ‘expressed ORF’ contigs (containing all contigs that mapped to a transcript, regardless of its inclusion on the genetic map; note that PAR contigs were included in the autosomal bin). There were 548 sex-linked contigs containing 1,641 genes and a total of 8.3 Mb of sequence; 7,579 autosomal contigs containing 23,502 genes and 106 Mb sequence data; and 15,392 contigs with 34,105 expressed ORFs distributed across 176 Mb (Table 1).

**Table 1.** Summary statistics of contigs containing sex-linked, autosomal or all expressed ORFs. Only ORFs supported by expression data were used for analysis of sex linkage (*de novo* predicted genes were excluded). Confidence intervals are one standard deviation.

|  | Whole genome | Expressed ORFs | Sex-linked | Autosomal |
| --- | --- | --- | --- | --- |
| Number of contigs | 720,537 | 15,392 | 538 | 7,579 |
| Total base pairs | 546,375,413 | 176,423,112 | 8,271,542 | 105,835,810 |
| % GC | 34.7 | 34.5 | 34.5 | 34.3 |
| Average contig length | 15,330 ($\pm 20,028$) | 11,462 ($\pm 17,207$) | 15,374 ($\pm 21,615$) | 13,964 ($\pm 19,678$) |
| Total repeat density (%) | 51.5 | 18.7 | 18.4 | 18.3 |
| Final coding transcript number | 34,105 | 34,105 | 1,641 | 23,502 |
| Average transcript length | 887 ($\pm 754$) | 810 ($\pm 702$) | 1,265 ($\pm 878$) | 1,242 ($\pm 853$) |
| Mean effective number of codons | 53.7 | 53.7 | 52.2 | 51.9 |
| Number of SNPs | 181,567 | 180,925 | 5,274 | 102,881 |
| Number of ORFs transcripts with SNPs | 16,667 (49%) | 16,572 (49%) | 545 (33%) | 7,714 (33%) |
| Median nucleotide diversity ($\pi$) / ORF | 0.0035 ($\pm 0.0113$) | 0.0035 ($\pm 0.0113$) | 0.006647 ($\pm 0.008079$) | 0.007951 ($\pm 0.01016$) |

### Sequence divergence, nucleotide diversity and codon usage of X and Y haplotypes

We used the SNP and haplotype calls from SEX-DETector to compare measures of nucleotide diversity and divergence between the X and Y ORFs within *M. annua*, as well as between X, Y and autosomal ORFs of *M. annua* and closely related species. The mean pairwise d_N_/d_S_ between X and Y haplotypes within *M.* annua was 0.182 (Table 2). For sex-linked transcripts, we excluded the few SNPs that did not show clear sex-linked segregation, but we could not apply such error correction for autosomes. Accordingly, π might be underestimated for sex-linked genes compared to autosomes (see Figure S3, Table 1), so that metrics associated with sex-linked sequences should be regarded as conservative.

**Table 2.** Average pairwise d_S_ and d_N_ for ORFs of *M. annua* from X/Y haplotype comparison within *M. annua* or against *M. huetii* and *R. communis* sequences. Autosomal ORF comparisons with the other species are also reported. Haplotypes with premature stop codons were excluded.

| | $d_S$ | $d_N$ | $d_N/d_S$ |
| --- | --- | --- | --- |
| X/Y pairs | 0.011 | 0.002 | 0.182 |
| X / *M. huetii* | 0.112 | 0.016 | 0.143 |
| Y / *M. huetii* | 0.113 | 0.017 | 0.150 |
| X / *R. communis* | 0.548 | 0.070 | 0.128 |
| Y / *R. communis* | 0.555 | 0.070 | 0.126 |
| *M. annua* / *M. huetii* | 0.106 | 0.014 | 0.149 |
| *M. annua* / *R. communis* | 0.565 | 0.079 | 0.140 |

The pairwise d_S_ and d_N_/d_S_ values for all possible orthologs between *M. annua* and its relatives *M. huetti* and *R. communis,* and the X and Y haplotypes, are presented in Table 2 and Figure 3. For autosomal genes, d_N_/d_S_ between *M. annua* and *M. huetii* and between *M. annua* and *R. communis*, respectively, was 0.149 and 0.140. Pairwise d_S_ was lower between X/Y gene pairs within *M. annua* than between orthologous autosomal genes in *M. annua* and *M. huetii* or *R. communis*.

**Figure 3:**
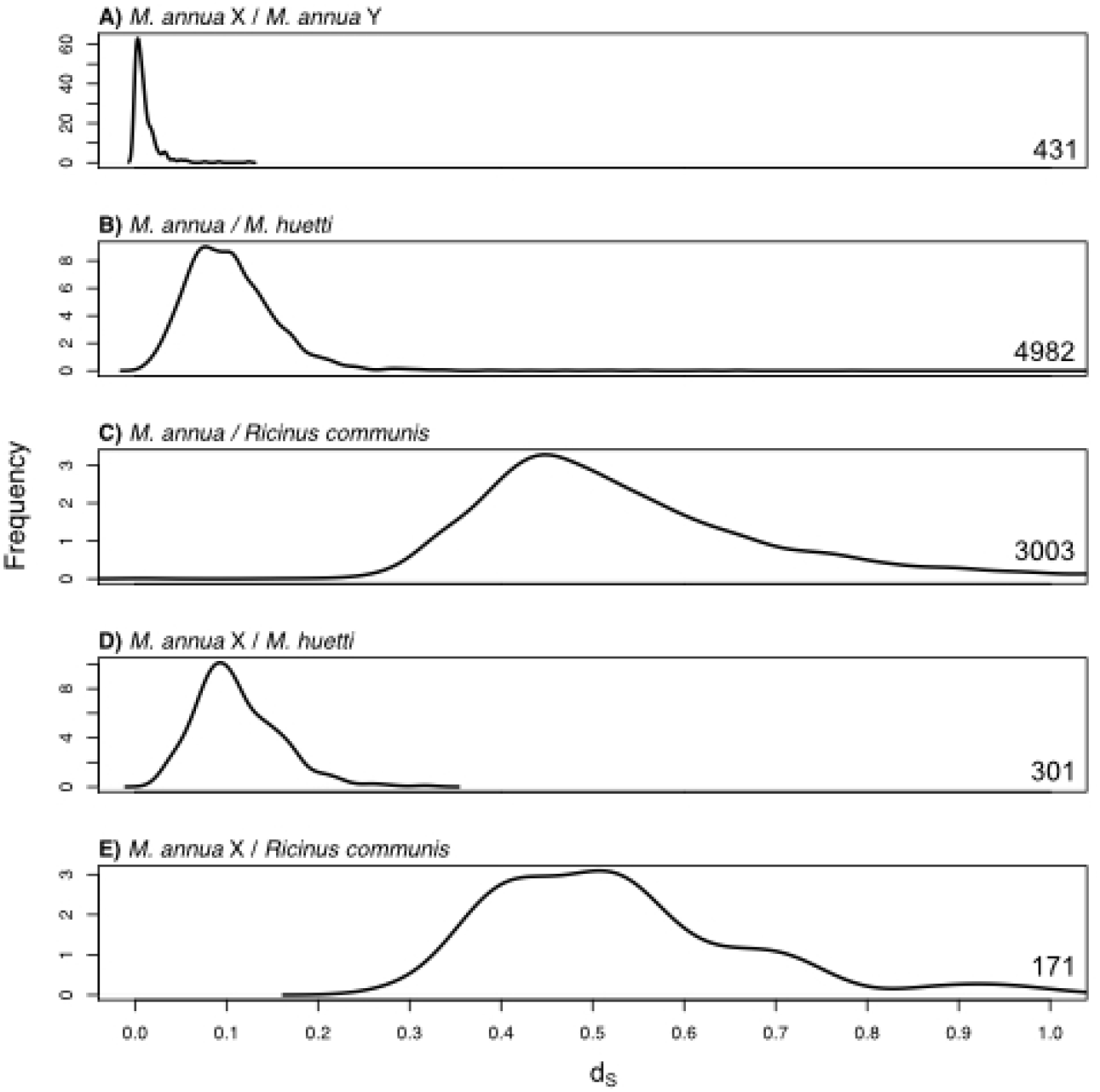
Density plots of synonymous site divergence (d_S_) for: (A) all *M. annua* X/Y haplotypes, (B) autosomal *M. annua* / *M. huetii* homologues, (C) autosomal *M. annua* / *R. communis* homologues, (D) X-linked *M. annua* / *M. huetii* homologues, and (E) X-linked *M. annua* / *R. communis* homologues. The numbers on the bottom right indicate the sample size for each plot.

Codon usage in *M. annua* did not differ significantly between sex-linked, pseudoautosomal and autosomal (SL, PAR and Au) ORFs (Nc = 54, Nc = 53 and Nc = 53, respectively; Figure S4).

We used PAML (Yang, 2007) to analyze the 98 sex-linked genes with orthologs in *M.annua, M. huetii*, and *R. communis*, using the ‘two-ratio’ M_2_model (which accommodates differences in d_N_/d_S_ between the Y branch and other branches of the tree; Nielsen and Yang 1998). Our analysis identified 84 sequences with 0.001 < d_S_ < 2 on both X and Y branches. Variation for 74 of these sequences was consistent with a tree topology corresponding to the species tree and recent X-Y divergence for *M. annua*, i.e. trees with topology (((*M. annua-*X, *M. annua-*Y), *M. huetii*), *R. communis*); we focused further analysis on these genes. For 52 of these 74 genes (21 after FDR correction at 5% level; Benjamini and Hochberg, 1995), the M_2_ model provided a better fit than the M_0_ model. Moreover, the Y lineage had more synonymous and more non-synonymous mutations compared to the X lineage, following divergence from *M. huetii*. Together, these results suggest that 28% of the Y-linked genes in our sample have been evolving faster than their corresponding X-linked orthologs as inferred from both tree-based d_N_ and d_S_ values (Wilcoxon Rank Sum = 392 and 568, respectively; p < 0.001 for both tests; Figure 4), as well as under relaxed selection as inferred from the tree-based d_N_/d_S_ ratio (Wilcoxon Rank Sum = 267, p < 0.001; Figure 4). The seven sequences for which the Y variant clustered outside the clade (*M. annua-*X, *M. huetii*) were distributed widely across genetic map of the non-recombining region of the Y (Figure 5), i.e. they did not clearly indicate a potential ancestral/common sex-determining region between *M. annua* and *M. huetii*. We did not investigate these genes further.

**Figure 4:**
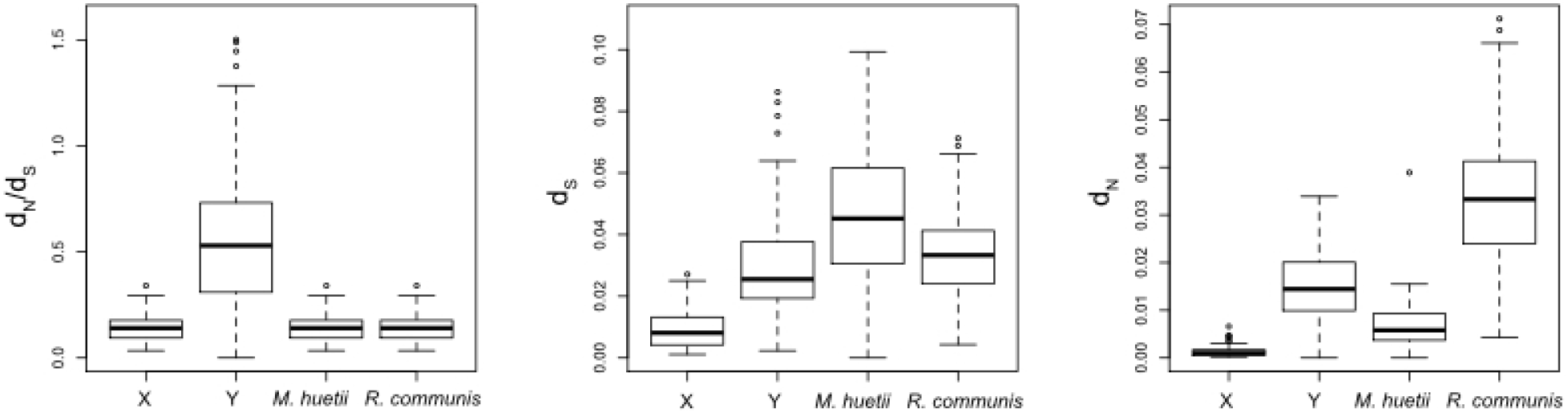
Boxplots of tree-based estimates of d_N_/d_S_, d_S_ and d_N_ for the X, Y, *M. huetii* and *R. communis* branches of the 72 gene trees that correspond to tree topology (((*M. annua-*X, *M. annua-*Y), *M. huetii*), *R. communis*). A Y-sequence d_N_/d_S_ outlier of 2.7 and an *M. huetii*-sequence d_S_ outlier of 0.23 are not shown, for greater resolution.

**Figure 5:**
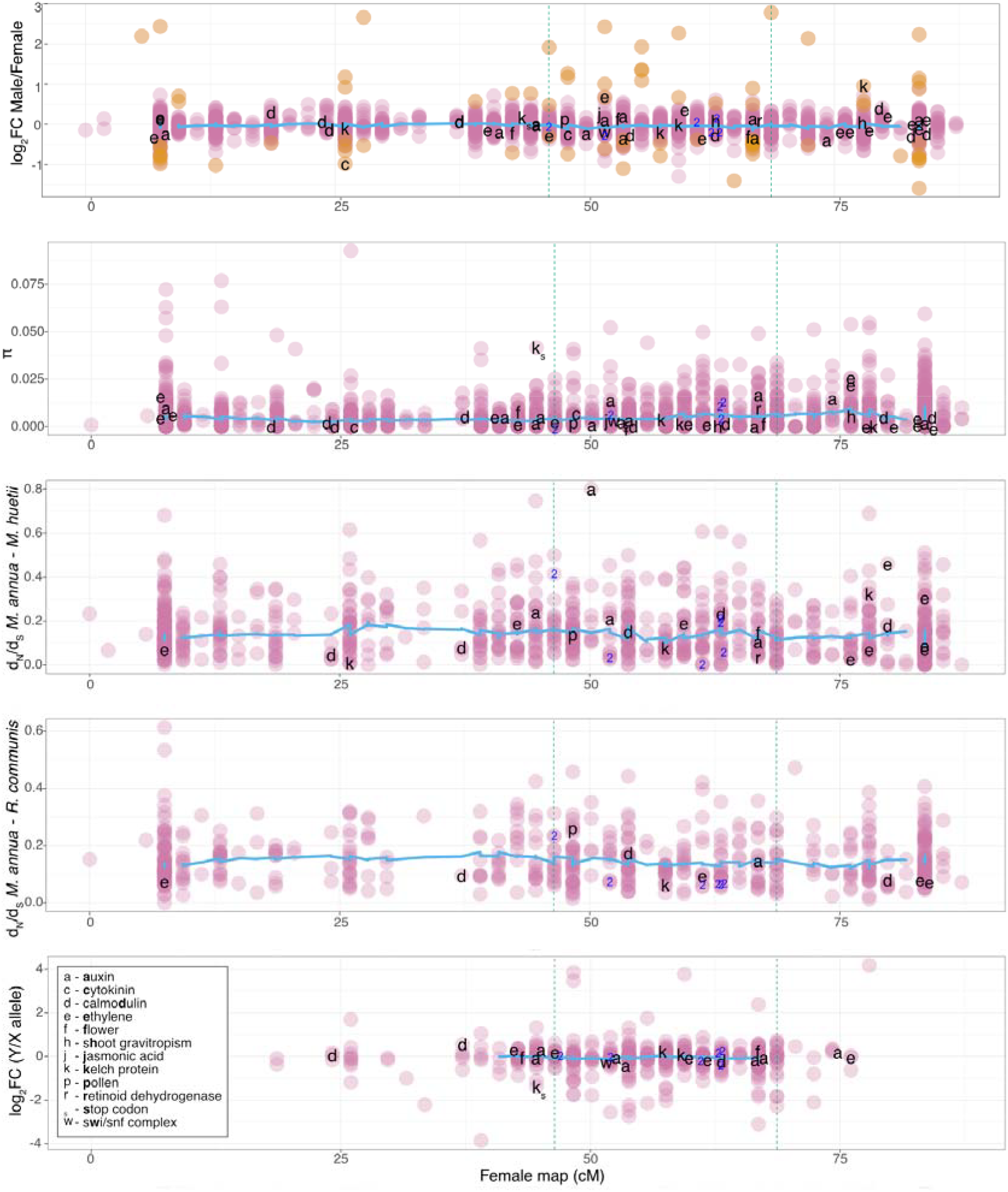
Summary of transcript metrics calculated for sequences across the female recombination map of the putative sex chromosome LG1, based on analysis of X and Y sequences combined. The non-recombining region in males is located between the vertical dotted lines. Orange points in the top panel indicate significant sex-bias (5% FDR). Potential candidate sex-determining genes are indicated by letters, with their putative function given in the inset legend. The only verified premature stop codon on the Y copy is indicated by k_s_ next to the non-recombining region (i.e., for a kelch protein gene). Transcripts where the Y sequence diverged before the *M. annua*/*M. huetii* species split are indicated with “2”.

### Identification of a degenerated and duplicated sequence on the Y

The SEX-DETector pipeline detected putative premature stop codons in four Y and three X transcripts, although there was always a functional X haplotype for the same transcript. We confirmed one of the Y haplotype stop codons using PCR, cloning and sequencing from five male and nine female samples from individuals sampled across the geographic range of diploid *M. annua*. Briefly, a phylogenetic tree was constructed for this gene using RAxML v 8.2.10 using 1,000 bootstraps (Stamatakis, 2014) on sequences aligned with mafft v7.310 (Katoh and Standley, 2013) (Figure S5). Interestingly, two different Y-linked sequences could be obtained, sometimes from the same male, suggesting that the sequence has duplicated in at least some males. All males and females analyzed also contained a functional copy which had 89% identity with an F-box/kelch repeat protein in *Ricinus communis*. The transcript shows lower Y/X expression ratio and high π (marked “K_s_” in Figure 5), suggesting that it represents a degenerated copy on the Y.

### Identification and annotation of sex-biased genes

Expression analysis revealed more genes with male-biased than with female-biased expression (1,385 male-biased vs. 325 female-biased genes, based on 5% FDR). The relative proportion of male-biased genes increased when also using a minimum log_2_fold-change (log_2_FC) threshold of one (1,141 vs. 140; Figure S6). An analysis of GO term enrichment is presented in File S3. Male-biased genes were enriched in biological functions such as anther-wall tapetum development, response to auxin, response to ethylene, cell-tip growth, pollen-tube growth, and floral-organ senescence. Female-biased genes were enriched for functions related to the maintenance of inflorescence meristem identity, jasmonic acid and ethylene-dependent systemic resistance, regulation of innate immune response, and seed maturation.

### Identification of the ancestral evolutionary stratum on the Y

To identify a putative ancestral stratum in the non-recombining region, we investigated variation in the following metrics across the female recombination map (Figure 5): the magnitude of sex-biased expression; the nucleotide diversity (π) of each transcript; and the pairwise d_N_/d_S_ ratio based on comparison with homologous genes in the closely related dioecious sister species *M. huetti*, and its monoecious more distant relative *R. communis* (for which we could identify 4,761 and 2,993 homologues, respectively); and the expression ratio of X and Y haplotypes inferred from SEX-DETector (Muyle *et al.*, 2016). We annotated Figure 5 with letters to indicate the position of candidate sex-determining genes, based on their blast hit descriptions. The sole obvious outlier for any of the metrics above was a transcript associated with auxin production, which had a high pairwise d_N_/d_S_ value in the comparison with the close outgroup species, *M. huetii*.

### Comparison of sex-linked, autosomal and pseudo-autosomal ORFs

We examined the effects of sex-bias (male-, female-, and unbiased genes) and genomic location (SL, PAR or Au), as well as their interactions, on the five metrics plotted in Figure 5. We repeated the analyses after excluding unbiased transcripts for the four metrics with sufficient gene numbers (sex bias log_2_FC, π, and pairise d_N_/d_S_ using either *M. huetii* or *R. communis*) to directly compare male-and female-biased transcripts. The full models are presented in Tables S9-S11. Below, we summarize their significant results; Figure 6 shows the data distributions. The analyses revealed significantly higher differences in gene expression between the sexes (absolute log_2_FC) in the sex-linked region (p = 0.037; Table S9). These differences were largely due to significantly higher expression of male-than female-biased transcripts, i.e. in terms of the extent of expression difference between the sexes, the sex chromosomes are significantly masculinized (male-bias*SL interaction: p=0.036; Table S10).

**Figure 6:**
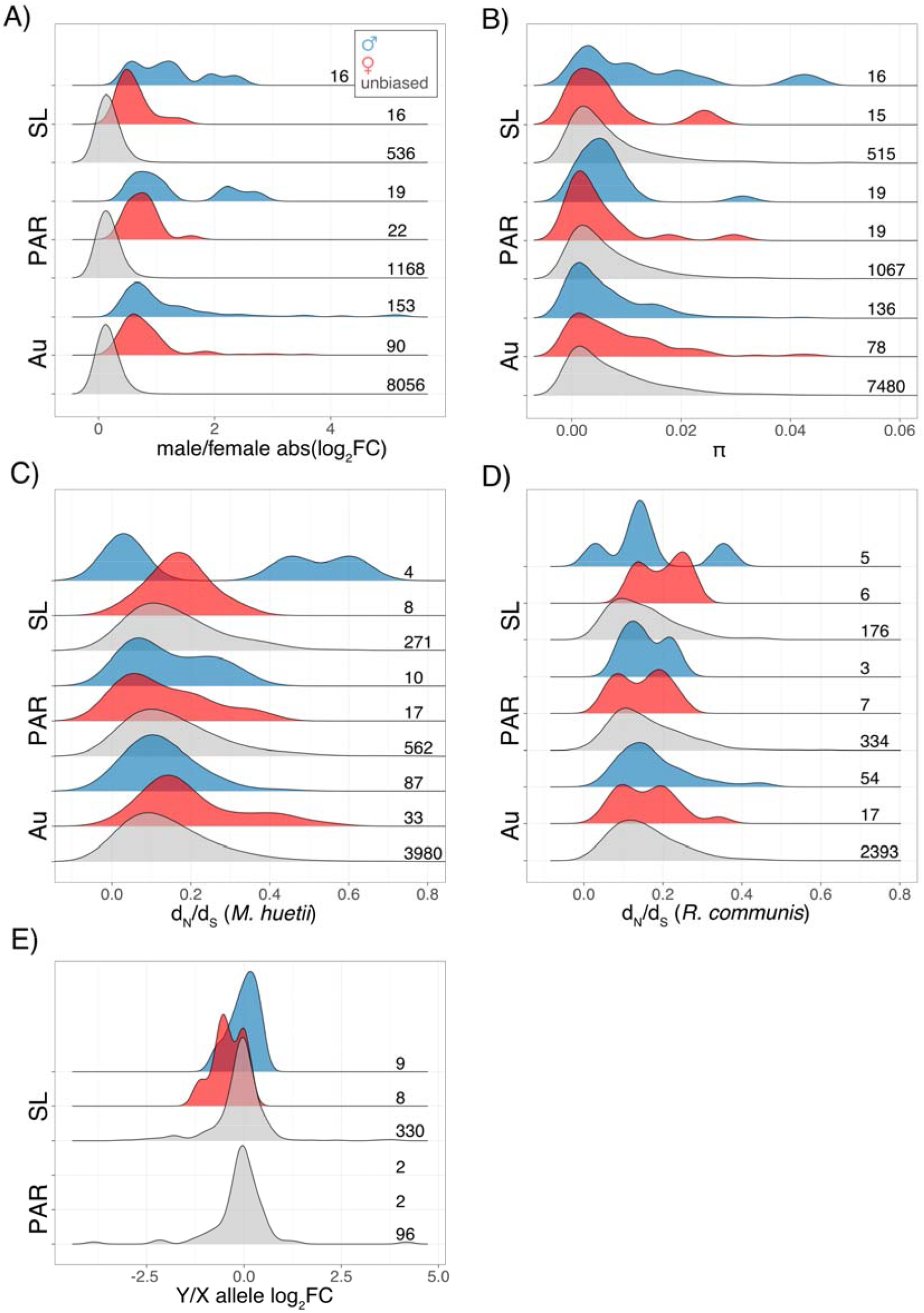
Graphical summary of the analysis of genes based on their sex-bias (blue: male-biased, red: female-biased, gray: unbiased) and sex linkage (SL: sex linked, PAR: remaining LG1, Au: LG 2-7). A) absolute log_2_FC expression difference between the sexes, B) nucleotide diversity (π), C) pairwise d_N_/d_S_ compared to *M. huetii* homologues, D) pairwise d_N_/d_S_ compared to *R. communis* homologues, E) relative expression of Y over X alleles. The numbers to the right indicate the number of genes in each category. Data are plotted prior to Box-Cox transformation.

The increased magnitude of male-biased gene expression in fully sex-linked genes is consistent with a recent masculinization as sex-linked genes had slightly lower nucleotide diversity (π) than those in the PAR or autosomal genes (p = 0.039; Table S9), but male-biased genes were not affected as much (p = 0.017; Table S9). The pairwise d_N_/d_S_ compared to *M. huetii* was higher for sex-linked genes (p = 0.017; Table S9) and for female-biased genes localizing to the PAR (PAR*female-bias interaction: p = 0.005; Table S9). An analysis of the sex-biased genes only (excluding unbiased) revealed the PAR to have a lower π (PAR: p = 0.007; Table S10) and a lower pairwise d_N_/d_S_ to *M. huetii* (p = 0.018; Table S10). Few patterns were significant for pairwise d_N_/d_S_ with respect to to the more distantly related *R. communis* (it was higher for male-biased genes; p = 0.046; Table S9), possibly because of lower power due to fewer identified homologues (4,761 in *M. huetti* compared to 2,993 in *R. communis*). The Y-linked copy of female-biased genes had significantly lower expression than the X-linked counterpart (p = 0.035; Table S11).

Finally, we used chi-squared tests to compare the relative proportions of sex-biased versus unbiased and male-versus female-biased genes in different pairs of genomic regions (PAR-SL, Au-LG1, Au-SL; Table 3). The sex-linked region had significantly more sex-biased genes (compared to either PAR or Au regions). This enrichment was largely responsible for the significant sex-biased gene enrichment compared with the entire LG1 and the autosomes, and was largely due to more female-biased genes on LG1. The sex chromosomes thus appear to be feminized in terms of number of female-biased genes, in contrast to the genome as a whole, which was masculinized in terms of number of male-biased gene numbers, and in contrast to absolute expression, which was higher for male-biased genes. We did not detect any enrichment for candidate genes with functions likely to have sexually antagonistic effects in the sex-linked region (Table 3).

**Table 3.** Results of chi-squared tests for differences in the frequencies of sex-biased versus -unbiased genes, male-versus female-biased genes, and candidate sex-determining genes (CG) versus other genes (not CG) between the pseudoautosomal region (PAR), the sex-linked region (SL), linkage group 1 (LG1) and autosomes (Au).

|  | Sex-biased vs unbiased |  |  | Female-biased vs male-biased |  |  | Candidate gene vs other gene |  |  |
| --- | --- | --- | --- | --- | --- | --- | --- | --- | --- |
|  | Unbiased | Sex-biased | Total | Female biased | Male biased | Total | Not CG | CG | Total |
| PAR | 1168 | 41 | 1209 | 22 | 19 | 41 | 1177 | 32 | 1209 |
|  | 96.6 % | 3.4 % | 100 % | 53.7 % | 46.3 % | 100 % | 97.4 % | 2.6 % | 100 % |
| SL | 536 | 32 | 568 | 16 | 16 | 32 | 546 | 22 | 568 |
|  | 94.4 % | 5.6 % | 100 % | 50 % | 50 % | 100 % | 96.1 % | 3.9 % | 100 % |
| Total | 1704 | 73 | 1777 | 38 | 35 | 73 | 1723 | 54 | 1777 |
|  | 95.9 % | 4.1 % | 100 % | 52.1 % | 47.9 % | 100 % | 97 % | 3 % | 100 % |
| Test | $\chi^2=4.381$ df=1 $\phi=0.053$<br>**p=0.036** | | | $\chi^2=0.006$ df=1 $\phi=0.036$<br>p=0.941 | | | $\chi^2=1.578$ df=1 $\phi=0.033$<br>p=0.209 | | |
| Au | 8056 | 243 | 8299 | 90 | 153 | 243 | 8094 | 207 | 8301 |
|  | 97.1 % | 2.9 % | 100 % | 37 % | 63 % | 100 % | 97.5 % | 2.5 % | 100 % |
| LG1 | 1704 | 73 | 1777 | 38 | 35 | 73 | 1723 | 54 | 1777 |
|  | 95.9 % | 4.1 % | 100 % | 52.1 % | 47.9 % | 100 % | 97 % | 3 % | 100 % |
| Total | 9760 | 316 | 10076 | 128 | 188 | 316 | 9817 | 261 | 10078 |
|  | 96.9 % | 3.1 % | 100 % | 40.5 % | 59.5 % | 100 % | 97.4 % | 2.6 % | 100 % |
| Test | $\chi^2=6.326$ df=1 $\phi=0.026$<br>**p=0.012** | | | $\chi^2=4.649$ df=1 $\phi=0.129$<br>**p=0.031** | | | $\chi^2=1.515$ df=1 $\phi=0.013$<br>p=0.218 | | |
| Au | 8056 | 243 | 8299 | 90 | 153 | 243 | 8094 | 207 | 8301 |
|  | 97.1 % | 2.9 % | 100 % | 37 % | 63 % | 100 % | 97.5 % | 2.5 % | 100 % |
| SL | 536 | 32 | 568 | 16 | 16 | 32 | 546 | 22 | 568 |
|  | 94.4 % | 5.6 % | 100 % | 50 % | 50 % | 100 % | 96.1 % | 3.9 % | 100 % |
| Total | 8592 | 275 | 8867 | 106 | 169 | 275 | 8640 | 229 | 8869 |
|  | 96.9 % | 3.1 % | 100 % | 38.5 % | 61.5 % | 100 % | 97.4 % | 2.6 % | 100 % |
| Test | $\chi^2=12.066$ df=1 $\phi=0.038$<br>**p<0.001** | | | $\chi^2=1.496$ df=1 $\phi=0.085$<br>p=0.221 | | | $\chi^2=3.493$ df=1 $\phi=0.021$<br>p=0.062 | | |

### Sex-linked sequence analysis of individuals sampled across the species range

To confirm the lack of recombination in the sex-linked region, we estimated *F*_ST_ and sex-biased heterozygosity for genome capture data from 20 males and 20 females sampled from natural populations across the species’ range (Table S2), based on 6,557 transcripts assigned to the linkage map. The sex-linked region showed clear differentiation between males and females using both metrics. *F*_ST_ between males and females was elevated in the sex-linked region relative to the PAR and the autosomes (Wilcoxon, p = 4.79×10^-5^, p = 1.099×10^-7^ respectively; Figure 7A), although the *F*_*ST*_ values are very small. Similarly, there was an excess of SNPs in males relative to females in the sex-linked region relative to the PAR and the autosomes (Wilcoxon, p<2.2×10^-16^ in both; Figure 7C). We used sliding window analyses of *F*_ST_ and SNP density to test whether males and females are more strongly differentiated in the sex-linked region compared to the rest of the genome. These analyses identified a region of elevated *F*_ST_ at ~ 65 cM of LG1 in the female recombination map (Figure 7B) whose value exceeded that for any autosomal region. We also identified a region between 50 and 65 cM on LG1 (Figure 7D) with higher SNP density and higher heterozygote frequencies in males than females (Figure 7E). Finally, we tested for enrichment for these genome-wide metrics from many populations. Table S12 shows that LG1 is both enriched in *F*_ST_ and sex-biased heterozygosity in the sex-linked region for males than females, as expected for X and Y chromosome divergence.

**Figure 7:**
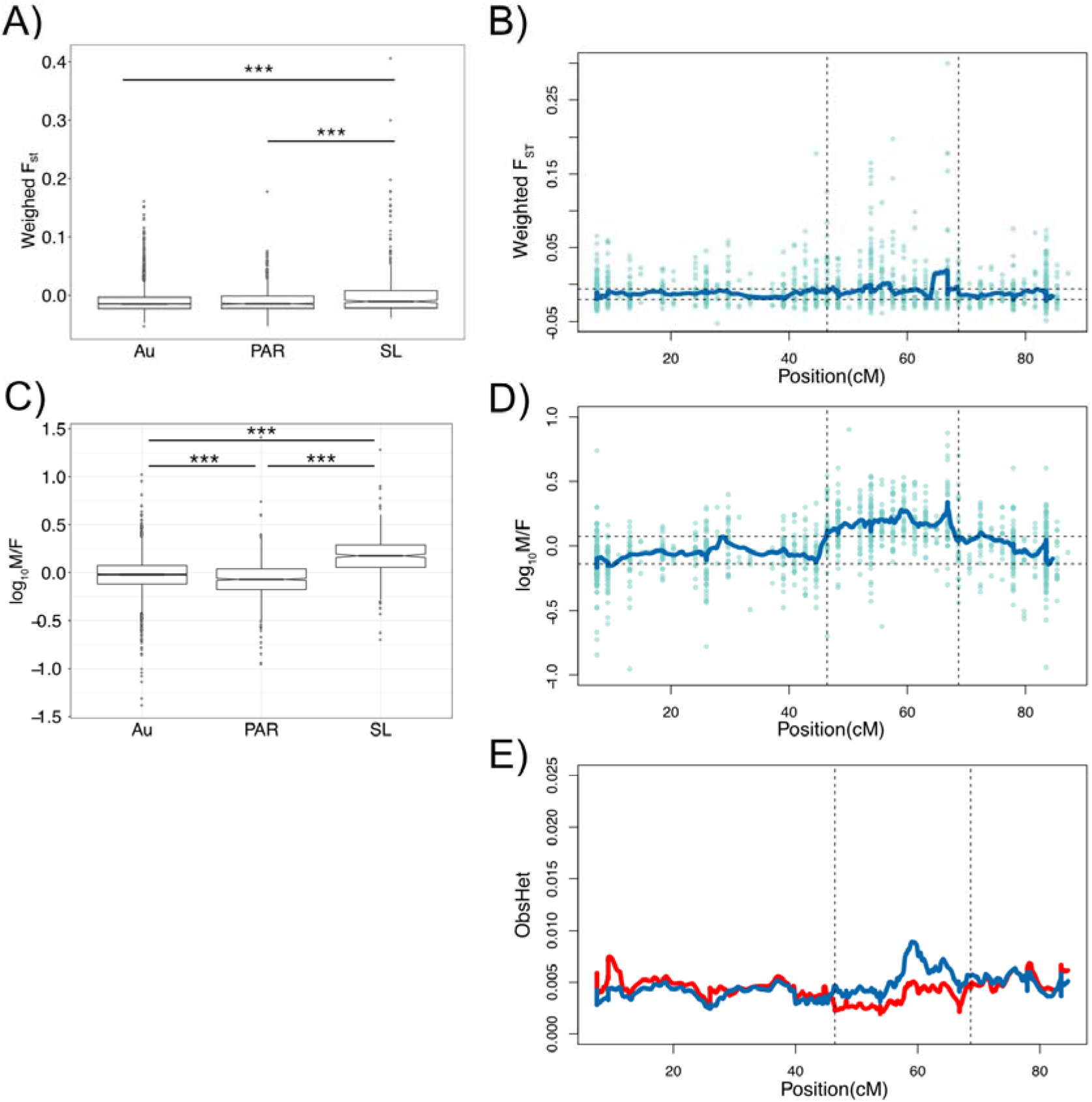
Summary of the effect of sex linkage on SNP metrics from genome capture data across the species range. (A) *F*_ST_ for sex linkage category, (B) *F*_ST_ distribution across the female recombination map of LG1, (C) ratio of male/female SNP density for sex linkage category, (D) ratio of male/female SNP density across the female recombination map of LG1, (E) observed heterozygosity in males (blue) and females (red) across LG1. Asterisks in A and C indicate significant Wilcoxon tests (p < 0.001), the sex-linked region is indicated by vertical lines in B, D and E.

## Discussion

### A genome assembly, annotation and genetic map for dioecious *M. annua*

Our study provides a draft assembly and annotation of diploid dioecious *M. annua* (2n = 16). Cytogenetic analysis has confirmed the haploid number of n = 8 chromosomes for diploid *M. annua* (Durand, 1963), with about 10,000 ORFs assigned to the corresponding eight linkage groups by genetic mapping. Similar to other plants, about 2/3 of the *M. annua* genome comprises repetitive sequences, mostly *gypsy* LTR, *copia* LTR, and L1 LINE retrotransposons (Chan *et al.*, 2010; Sato *et al.*, 2011; Wang *et al.*, 2012a; Rahman *et al.*, 2013). The number of genes in *M. annua* (34,105) is more similar to that of diploid species such as *Arabidopsis thaliana* (> 27,000 genes) and *Ricinus communis* (> 31,000 genes) and is thus compatible with having a long recent history under diploidy rather than polyploidy; for comparison, the gene content of diploid *Gossypium* is > 40,000 genes, pointing to a polyploid history (Wang *et al.*, 2012a). The draft annotated genome of *M. annua* will be useful for understanding potential further links between sex determination and sexual dimorphism, as well as the consequences of genome duplication and hybridization (Obbard *et al.*, 2006) for sexual-system transitions in polyploid lineage of the species complex (Pannell *et al.*, 2008).

### The size of the non-recombining Y-linked region of *M. annua*

Patterns of SNP segregation in small families from 568 ORFs suggests that a substantial length (1/3) of the *M. annua* Y chromosome has ceased recombining with the X. Lack of recombination on the sex-linked region is further supported by the existence of a Y-specific non-functional gene duplication, which could be amplified in populations ranging from Israel to Britain, as well as additional population-genetic data from males and females sampled from across the diploid species’ range. The sex-linked region had slightly, but significantly, higher *F*_ST_, a higher male/female SNP ratio, and higher heterozygote frequency in males than females, indicating a history of low recombination. Previous mapping of bacterial artificial chromosomes (BACs) containing male-specific PCR products found them to be distributed over most of the sex-linked region, i.e., a region of between 52 and 66.82 cM, corresponding to 4.86% of the genome and implying a physical length of between 14.5 and 19 Mb for the non-recombining genome (Veltsos *et al.*, 2018). The large non-recombining region of the *M. annua* Y chromosome contrasts with that of other plants with homomorphic sex chromosomes, which typically have < 1% of their Y not recombining, e.g., *Vitis vinifera* (Fechter *et al.*, 2012; Picq *et al.*, 2014), *Fragaria chiloensis* (Tennessen *et al.*, 2016) and *Populus* species (Paolucci *et al.*, 2010; Geraldes *et al.*, 2015).

### Limited gene loss, pseudogenization, and mild purifying selection on the *M. annua* Y

We found only limited evidence for Y-chromosome degeneration within the non-recombining region of the *M. annua* Y chromosome, despite its substantial length. Very few genes have been lost or have become non-functional, and only one of the 528 X-linked genes corresponding to the non-recombining region of the Y (0.2%) did not have a Y-linked homologue. Our analysis may have overlooked genes that have been lost from the Y, because their detection from RNAseq data relies on the presence of polymorphisms on the X copy (see Blavet *et al.*, 2015) and requires their detectable expression in our sampled tissues. Nevertheless the absence of evidence for substantial gene loss is consistent with other signatures of only mild Y-chromosome degeneration in *M. annua* and is likely a reliable finding.

There was only slightly lower relative expression from the Y compared to the X allele. Allele-specific expression was inferred on the basis of mapping to the reference transcriptome, and we expect only limited mapping bias against expressed Y-specific genes. The fact that different levels of Y/X expression depended on sex-bias (it was reduced for female-biased genes) further suggests that the result is probably not an artifact of the analysis. We found some evidence for relaxed purifying selection on Y alleles compared to X alleles, as expected in genomic regions experiencing limited recombination (Charlesworth and Charlesworth, 2000). Specifically, pairwise comparisons of X and Y alleles (after excluding the few with in-frame stop codons) showed higher d_N_/d_S_ than pairwise comparisons of *M. annua* orthologs with *M. huetii* or *R. communis*. Moreover, our tree-based analysis of common orthologs in all species indicated that 28% of the identified Y-linked alleles have experienced an accelerated rate of molecular evolution in the *M. annua* lineage since its divergence from its sister species *M. huetii*, which is also dioecious. However, we found no evidence for differential codon usage bias between the Y-linked and other sequences. Codon usage bias is expected to be lower for non-recombining regions of the genome experiencing weaker purifying selection (Hill and Robertson, 1966), and there appears to have been a shift towards less preferred codon usage in *Rumex hastatulus* Y-linked genes, increasing in severity with time since the putative cessation of recombination between X and Y chromosomes (Hough *et al.*, 2014). Although our analysis involves multiple statistical tests of significance such that some results might represent type 1 error, taken together they point to a very recent cessation of recombination for much of the Y chromosome. In contrast, up to 28% of *Rumex hastatulus* Y-linked genes have been lost (Hough *et al.*, 2014) and about 50% of the *Silene latifolia* Y-linked genes are dysfunctional (Papadopulos *et al.*, 2015; Krasovec *et al.*, 2018).

### Sex-biased gene expression in *M. annua*

We found substantial differences in gene expression between males and females of *M. annua*, consistent with the observation of sexual dimorphism for a number of morphological, life-history and defense traits in *M. annua* (Hesse and Pannell, 2011; Labouche and Pannell, 2016). There were at least four times more male-biased than female-biased genes; this is probably conservative because we did not use a minimum log_2_FC threshold in our analysis, which would have further increased the proportion of male-biased genes. The observed slight decrease in expression of Y-compared with X-linked alleles suggests that dosage compensation could have begun evolving. However, any such dosage compensation would likely be at a very early stage, not least because YY males (which lack an X chromosome) are known to be viable (Kuhn, 1939). Interestingly, sex-linkage sometimes influenced male-and female-biased genes differently, with male-biased genes having a higher nucleotide diversity (<u>π)</u> than female-biased genes when sex-linked, again perhaps reflecting a history of relaxed purifying selection. Male-and female-biased genes also differed in pairwise d_N_/d_S_ relative to *M. huetii*, which was higher for female-biased genes, contrary to expectations under a hypothesis of faster male evolution (Wu and Davis, 1993).

### Evidence for evolutionary strata on the *M. annua* Y?

We were unable to find any direct evidence for evolutionary strata with different levels of divergence on the *M. annua* Y chromosome, but phylogenetic comparisons point to the likely existence of at least two such strata on the *M. annua* Y. Dioecy evolved in a common ancestor to *M. annua* and *M. huetii* (Krähenbühl *et al.*, 2002; Obbard *et al.*, 2006), and both species share the same Y chromosome and sex-determining region, as revealed by crosses (Russell and Pannell, 2015) and the possession of common male-specific PCR markers (Veltsos *et al.*, 2018). The phylogeny of sex-linked genes within the shared sex-determining region should have the topology ((*M. annua*-Y, *M. huetii*), *M. annua-*X). In contrast, our PAML analysis clearly indicates that the phylogeny of most genes in the non-recombining region of the *M. annua* Y have the topology ((*M. annua*-Y, *M. annua*-X), *M. huetii*), indicating divergence between the X and Y copies of these genes in *M. annua* more recently than the species split between *M. annua* and *M. huetii*. These results suggest that *M. annua* and *M. huetii* share an old (and probably small) non-recombining stratum that includes the sex-determining locus, and that a larger stratum has been added to the non-recombining region in the *M. annua* lineage. Another possible interpretation is gene conversion between X and Y copies within *M. annua*, as has been reported for humans and some plants (Rautenberg *et al.*, 2008; Wu *et al.*, 2010; Trombetta *et al.*, 2014). This would lead to an underestimate of the size of the ancestral stratum and may have erased evidence of a longer common shared history between the non-recombining regions of *M. annua* and *M. huetii*.

Although most Y-linked genes that we sampled in *M. annua* point to an X/Y divergence in *M. annua* that postdates the split from *M. huetii*, we found seven genes with a phylogeny consistent with the hypothesized old stratum shared with *M. huetii*. If these genes are indeed within an older stratum, we might expect them to colocalize in the same part of the non-recombining region. However, we found that they were quite scattered on the female recombination map. We have no explanation for this pattern, but we note that it would be consistent with a break of synteny between the *M. annua* Y and X chromosomes (and thus potentially also between the *M. huetii* and *M. annua* Y chromosomes) in the region corresponding to the non-recombining region.

Patterns of divergence in terms of tree-based d_S_ and d_N_ between the X and Y chromosomes of *M. annua* and their corresponding orthologs in *M. huetii* and *R. communis* were largely consistent with the expected phylogenetic relationships between these species (Krähenbühl *et al.*, 2002; Obbard *et al.*, 2006). The increased d_S_ for the Y branch is consistent with a higher mutation rate in male meiosis. Nevertheless, the d_S_ branch length for *R. communis* was lower than expected for a simple molecular clock across the phylogeny. The recent transition to an annual life cycle in the clade that includes *M. annua* and *M. huetii* may be partially responsible for the larger number of mutations that have accumulated along their branches compared to the perennial *R. communis* (Smith and Donoghue, 2008).

### Age of the non-recombining Y-linked region of *M. annua*

We may estimate the age of the *M. annua* non-recombining region on the basis of inferred mutation rates in plants. Ossowski *et al.* (2010) estimated the mutation rate for *Arabidopsis thaliana* as between 7 × 10^−9^ to 2.2 × 10^−8^ per nucleotide per generation. More recently, Krasovec *et al.* (2018) estimated the mutation rate for *S. latifolia* as 7.31 x 10^-9^. The similarity of these estimates suggests that they might be broadly applicable. Accepting a mutation rate for plants of approximately 7.5 × 10^−9^ and using the synonymous site divergence between X-and Y-linked sequences of *M. annua* of 0.11 (Table 2), we infer that recombination between most of the X-and Y-linked genes in *M. annua* ceased as recently as 1.5 million generations ago (or even more recently if we adopt the upper end of the mutation-rate range estimated by Ossowski *et al.* (2010). Given that *M. annua* is an annual plant, the putative second non-recombining stratum of the Y chromosome of diploid *M. annua* could be less than a million years old. Krasovec *et al.* (2018) used their mutation-rate estimate to infer an age of 11 million years for the oldest sex-linked stratum of *S. latifolia*, i.e., an order of magnitude older than the expanded sex-linked region of *M. annua*. The few estimates for the time since recombination suppression in other plants with homomorphic sex chromosomes, and those with smaller non-recombining regions, range from 15 to 31.4 million years ago (reviewed in Charlesworth, 2016). *Mercurialis annua* thus appears to have particularly young sex chromosomes.

### Concluding remarks

We conclude by speculating on a possible role for sexually antagonistic selection in favoring suppressed recombination on the *M. annua* Y. The canonical model for sex-chromosome evolution supposes that suppressed recombination originally evolves in response to selection to bring sexually antagonistic loci into linkage with the sex-determining locus, and that the Y chromosome begins to degenerate after recombination has ceased (Rice, 1987). Given this model, we might expect to see signatures of sexual antagonism in the sex-linked region before the onset of substantial Y-chromosome degeneration. While we do not observe obvious signatures of sexually antagonistic selection, some of the observed patterns in the sex-linked region are difficult to interpret as the outcome of degeneration. For example, while the Y-linked alleles showed slightly lower expression than X-linked alleles, this was not true for male-biased genes. One possible explanation is that the female-biased Y-linked alleles in males might have been driven to lower expression by sexually antagonistic selection. If female-biased expression is beneficial in females (Parisi *et al.*, 2003; Connallon and Clark, 2010), the observed enrichment of female-biased genes on the X would be consistent with sexually antagonistic selection because X-linked alleles occur twice as often in females than in males. Similarly, the magnitude of sex-biased gene expression (absolute log_2_FC) was higher in the sex-linked region, as might be expected if levels of gene expression were the result of an arms race between the sexes.

Our study prompts investigations in several directions. The speculation that sexually antagonistic selection might have played a role in the evolution of suppressed recombination might be investigated further by seeking sex-linked QTL for sexually antagonistic traits (Delph *et al.*, 2010), or functional analysis of candidate genes. It would also be valuable to compare expression levels of sex-linked genes with those in a related species with ancestral levels of gene expression (see Zemp *et al.*, 2016). The orthologs of the *M. huetii* genes that are non-recombining only in *M. annua* would make for a good comparison. More generally, the *M. annua* species complex offers itself as a promising system for comparative analysis of diverging sex chromosomes within a small clade in which speciation, hybridization and genome duplication have contributed to diversification. We anticipate characterizing the sex-linked region of *M. huetii*, the sister species of *M. annua*, and to trace sex determination and the sex chromosomes from diploid *M. annua* into the related polyploid lineages that appear to have lost and then regained dioecy (Pannell *et al.*, 2004; Obbard *et al.*, 2006). These other lineages have very similar morphological differences between males and females, but may determine sex using different loci or have sex chromosomes with a different history.

## Supporting information

File S1 - Mapped transcripts

File S2 - tree stop codons

File S3 - GO results

File S4 - Transcriptome Annotation

File S5 - SD transcriptome

File S6 - genome

File S7 - Mercurialis repeat library

## Acknowledgements

We thank John Russell for setting up the crosses, Jos Käfer for help with the Galaxy workflow for the SEX-DETector analysis, Jerome Goudet for statistical advice, and Guillaume Cossard, Wen-Juan Ma, Deborah Charlesworth, Stephen Wright and anonymous reviewers for valuable comments on the manuscript. KER was supported by grants 31003A_163384 and 31003A_141052 to JRP from the Swiss National Science Foundation, and by the University of Lausanne. PV was supported by Sinergia grant 26073998 from the Swiss National Science Foundation to JRP, Nicolas Perrin and Mark Kirkpatrick. JRP and DAF acknowledge support from the NERC and BBSRC, UK, which funded the early stages of this project. The computations were performed at the server at the department of Plant Sciences, University of Oxford, the Vital-IT (http://www.vital-it.ch) Center for high-performance computing of the SIB Swiss Institute of Bioinformatics and the Texas Advanced Computing Center. AM and GABM acknowledge support from ANR (grant number ANR-14-CE19-0021-01). VH, RH, BV were supported by the Czech Science Foundation (grant 18-06147S). MAT was supported by the National Institute of Health grant R01GM116853 (to Mark Kirkpatrick) and the European Research Council under the European Union’s Horizon 2020 research and innovation program (grant agreement number 715257).

## Author contributions

Conceived the project: JRP, DAF; wet lab work: DAF; cytogenetics work: VH, RH, PV; genome assembly and annotation: KER; gene expression analysis: KER, OE, PV; segregation analysis and mapping: PV, AM, PR, GM; population genetic and divergence analysis: KER, PV; analysis of natural populations: SCGM, MAT: wrote the paper: JRP, KER, PV; commented and contributed to the final manuscript: all authors.

## Supplementary Information: Tables

**Table S1:**
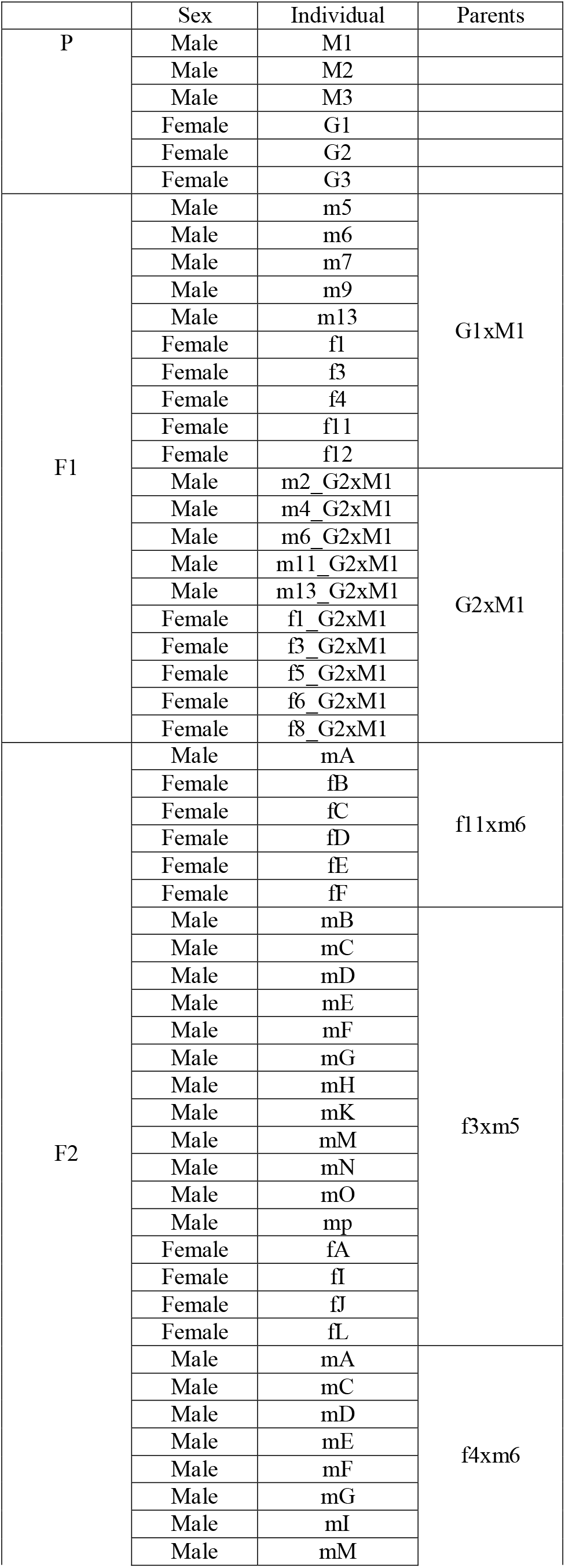
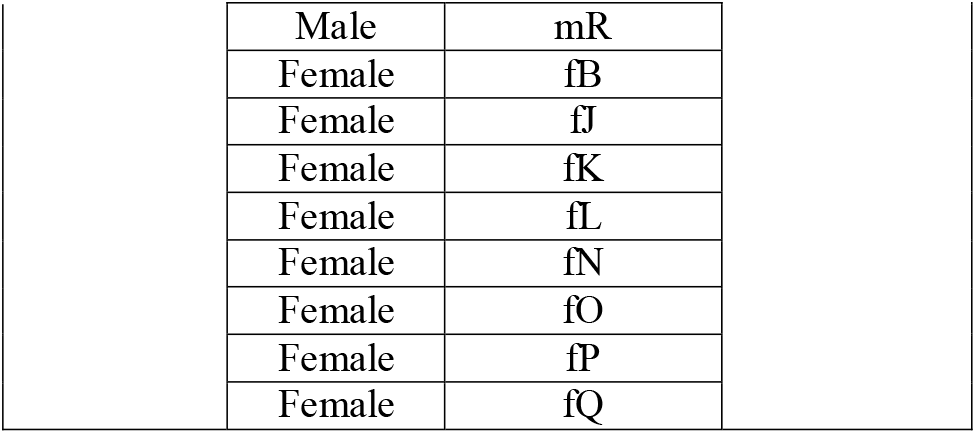
Samples from wild accessions and genetic cross progeny used for RNAseq.

|  | Sex | Individual | Parents |
| --- | --- | --- | --- |
| P | Male | M1 |  |
|  | Male | M2 |  |
|  | Male | M3 |  |
|  | Female | G1 |  |
|  | Female | G2 |  |
|  | Female | G3 |  |
| F1 | Male | m5 | G1xM1 |
|  | Male | m6 |  |
|  | Male | m7 |  |
|  | Male | m9 |  |
|  | Male | m13 |  |
|  | Female | f1 |  |
|  | Female | f3 |  |
|  | Female | f4 |  |
|  | Female | f11 |  |
|  | Female | f12 |  |
|  | Male | m2_G2xM1 | G2xM1 |
|  | Male | m4_G2xM1 |  |
|  | Male | m6_G2xM1 |  |
|  | Male | m11_G2xM1 |  |
|  | Male | m13_G2xM1 |  |
|  | Female | f1_G2xM1 |  |
|  | Female | f3_G2xM1 |  |
|  | Female | f5_G2xM1 |  |
|  | Female | f6_G2xM1 |  |
|  | Female | f8_G2xM1 |  |
| F2 | Male | mA | f11xm6 |
|  | Female | fB |  |
|  | Female | fC |  |
|  | Female | fD |  |
|  | Female | fE |  |
|  | Female | fF |  |
|  | Male | mB | f3xm5 |
|  | Male | mC |  |
|  | Male | mD |  |
|  | Male | mE |  |
|  | Male | mF |  |
|  | Male | mG |  |
|  | Male | mH |  |
|  | Male | mK |  |
|  | Male | mM |  |
|  | Male | mN |  |
|  | Male | mO |  |
|  | Male | mp |  |
|  | Female | fA |  |
|  | Female | fI |  |
|  | Female | fJ |  |
|  | Female | fL |  |
|  | Male | mA | f4xm6 |
|  | Male | mC |  |
|  | Male | mD |  |
|  | Male | mE |  |
|  | Male | mF |  |
|  | Male | mG |  |
|  | Male | mI |  |
|  | Male | mM |  |

|  |  |
| --- | --- |
| Male | mR |
| Female | fB |
| Female | fJ |
| Female | fK |
| Female | fL |
| Female | fN |
| Female | fO |
| Female | fP |
| Female | fQ |

**Table S2:**
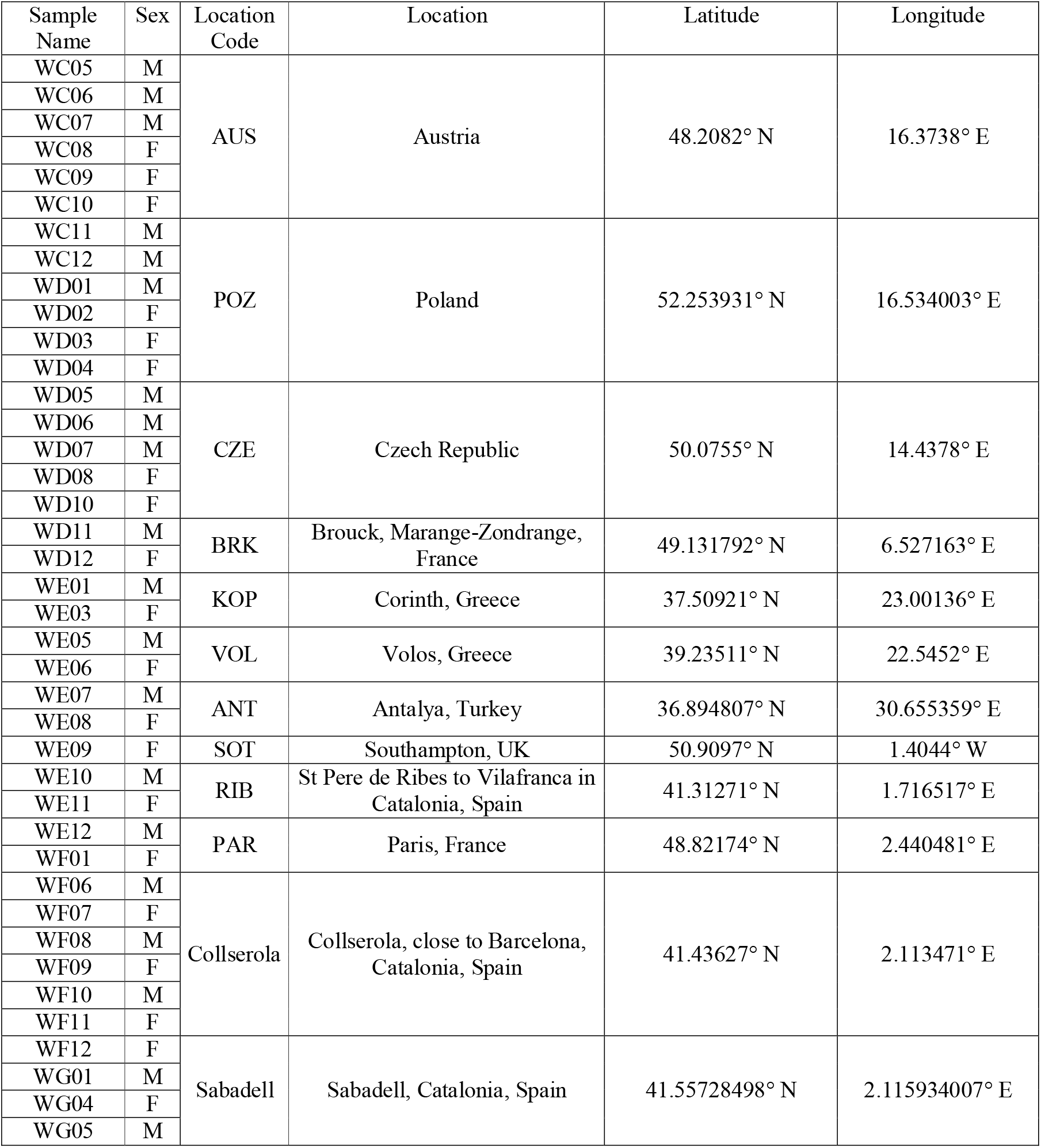
Samples from wild accessions used for genome capture data.

**Table S3:** Summary of arm ratio (r=p/q) and relative chromosome length ((p+q)/absolute length * 100) based on 50 figures from 5 individuals. All measurements were done as in Lengerova *et al.* (2004).

| Chromosome number | Arm ratio (r) | SD | Relative length |
| --- | --- | --- | --- |
| 1 | 1.13 | $\pm 0.05$ | 20.17 |
| 2 | 1.93 | $\pm 0.12$ | 14.56 |
| 3 | 2.36 | $\pm 0.40$ | 14.53 |
| 4 | 1.53 | $\pm 0.05$ | 11.32 |
| 5 | 1.31 | $\pm 0.12$ | 10.64 |
| 6 | 1.25 | $\pm 0.15$ | 9.23 |
| 7 | 0.96 | $\pm 0.00$ | 10.24 |
| 8 | 1.14 | $\pm 0.24$ | 9.31 |

**Table S4:** Summary results from k-mer (size 31) analysis.

|  | M1 |  | G1 |  | G2 |  |
| --- | --- | --- | --- | --- | --- | --- |
|  | min | max | min | max | min | max |
|  | 1.85% | 1.88% | 1.82% | 1.85% | 1.83% | 1.86% |
| Haploid length (Mb) | 325.20 | 327.06 | 292.72 | 294.39 | 344.57 | 346.80 |
| Repeat length (Mb) | 126.45 | 127.17 | 101.09 | 101.67 | 142.62 | 143.55 |
| Unique length (Mb) | 198.75 | 199.89 | 191.63 | 192.72 | 201.95 | 203.26 |
| Model fit | 92.41% | 98.17% | 92.77% | 97.68% | 91.99% | 98.27% |
| Read error rate | 0.47% | 0.47% | 0.49% | 0.49% | 0.35% | 0.35% |

**Table S5:**
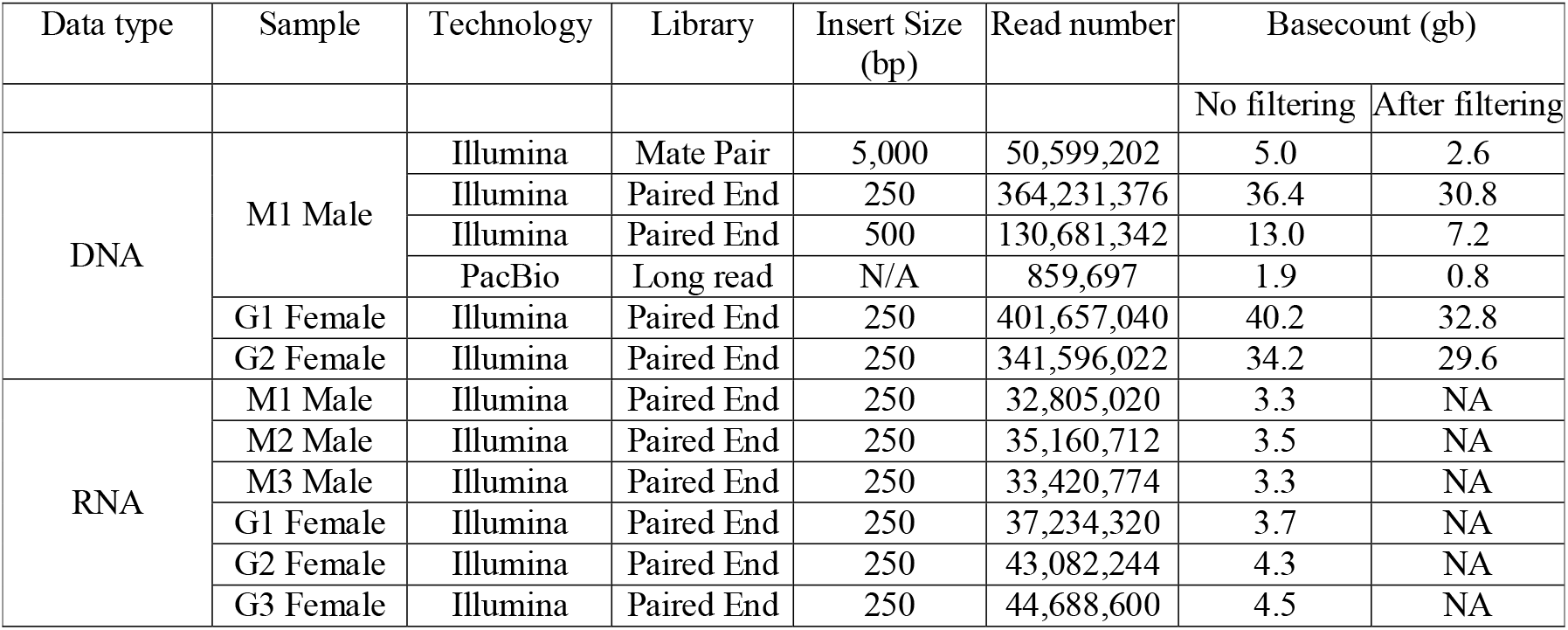
Raw data before and after filtering of genomic and transcript libraries. RNA-seq was not filtered, as those reads would drop out during the assembly and ORF prediction.

**Table S6:** Summary of the genome *de novo* assembly. Scaffold lengths are displayed in base pairs. Software was run sequentially from SOAP2 to L_RNA_scaffolder.

| Software | Scaffold number | Total bases (% coverage) | Bases without gaps (% coverage) | Longest scaffold | Scaffold n50 |
| --- | --- | --- | --- | --- | --- |
| SOAP2+GapCloser | 832,132 | 526,869,874 (86%) | 455,924,417 (74%) | 335,033 | 5,903 |
| PBJelly2 | 825,889 | 547,240,256 (89%) | 481,594,113 (78%) | 329,358 | 5,642 |
| SSPACE | 726,400 | 545,659,344 (89%) | 479,897,493 (78%) | 329,358 | 6,126 |
| LRNA scaffolder | 720,537 | 546,375,413 (89%) | 479,897,493 (78%) | 329,358 | 6,398 |
| Final over 1kb | 74,927 | 397,889,956 (65%) | 332,624,619 (54%) | 329,358 | 12,808 |

**Table S7:** The *de novo* assembly of *M. annua* compared to the assemblies of the other sequenced members of the Malpighiales.

|  | Genome size<br>(mb) | Scaffold number | Total assembled<br>bases | Scaffold n50<br>(bp) | Citation |
| --- | --- | --- | --- | --- | --- |
| *Mercurialis annua*<br>(Annual mercury) | 640 | 720,537 | 546,375,413 (85%) | 6,398 | (Obbard *et al.*, 2006;<br>this study) |
| *Ricinus communis*<br>(Castor bean) | 400 | 25,763 | 350,621,860 (88%) | 496,528 | (Chan *et al.*, 2010) |
| *Populus trichocarpa*<br>(Poplar) | 485 | 2,514 | 417,286,671 (86%) | 18,835,763 | (Tuskan *et al.*, 2006) |
| *Linum usitatissimum*<br>(Flax) | 350 | 88,420 | 318,250,901 (91%) | 693,500 | (Wang *et al.*, 2012c) |
| *Manihot esculenta*<br>(Cassava) | 760 | 12,977 | 416,657,463 (55%) | 180,909 | (Prochnik *et al.*, 2012) |
| *Hevea brasiliensis*<br>(Rubber tree) | 2,000 | 608,017 | 1,100,000,000<br>(55%) | 2,972 | (Tang *et al.*, 2016) |
| *Jatropha curcas*<br>(Oil-bearing tree) | 410 | 39,277 | 297,660,620 (73%) | 15,950 | (Sato *et al.*, 2011) |
| *Salix suchowensis*<br>(Shrub willow) | 429 | 103,144 | 303,767,511 (71%) | 924,964 | (Dai *et al.*, 2014) |

**Table S8:**
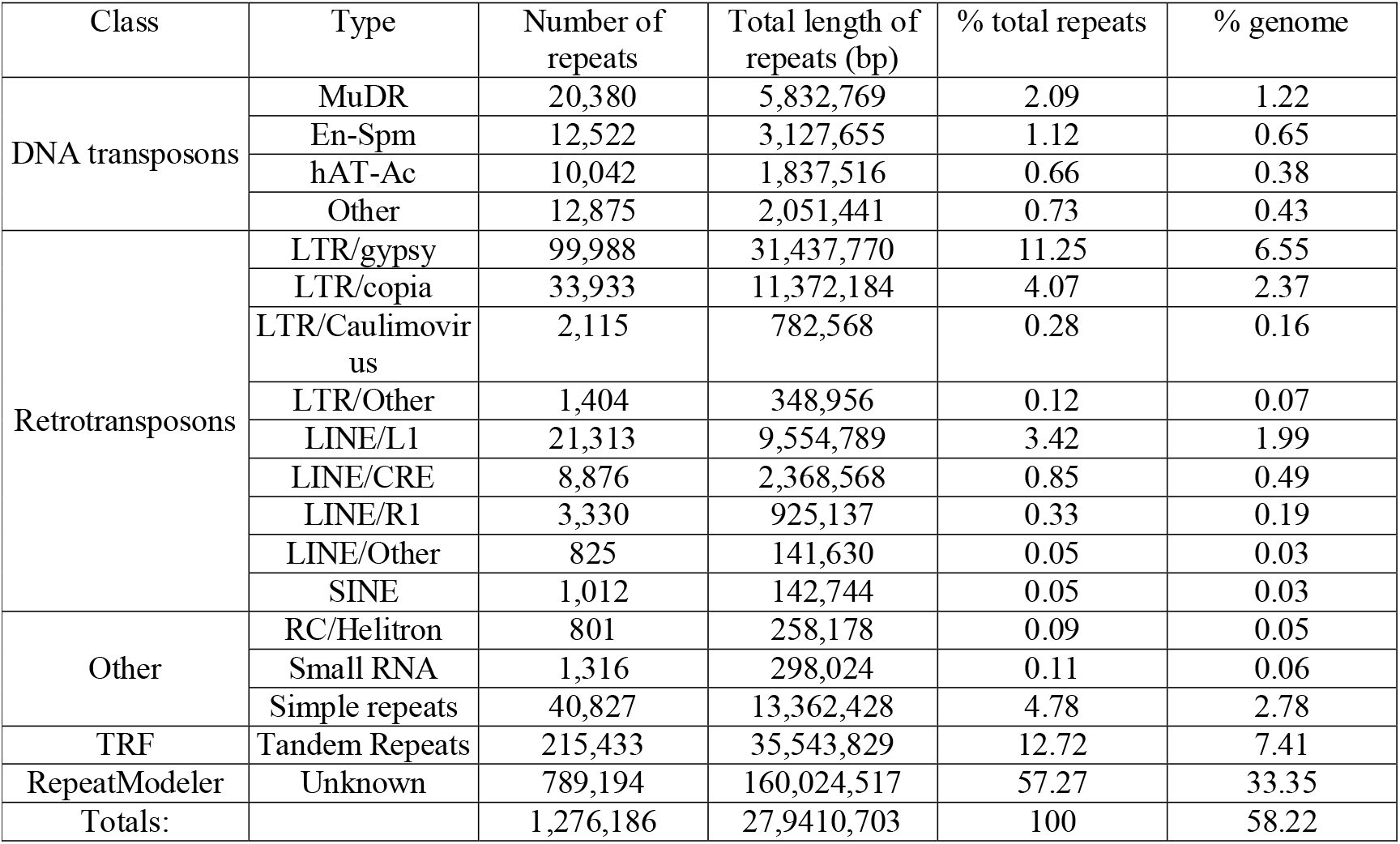
Transposable elements in the *Mercurialis* genome. Tandem repeats were identified using Tandem Repeats Finder (TRF). Unknown transposable elements were identified using RepeatModeler for *de novo* transposable element prediction. All other repeats were predicted using Repeat Masker, and repeats from *Euphorbiacae* and *Vitis vinifera*.

**Table S9:** Linear models on Box-Cox-transformed measures of sex-biased gene expression across all genes in *M. annua* (absolute log_2_FC difference between the sexes), nucleotide diversity (π), and pairwise dN/dS (with reference to *M. huetii* and *Ricinus communis)*, as a function of sex linkage (SL: sex-linked; PAR: pseudoautosomal, Au: autosomal), sex-bias (male: male-biased; female: female-biased) and their interactions.

| Predictors |  | Dependent Variables |  |  |  |  |  |  |  |
| --- | --- | --- | --- | --- | --- | --- | --- | --- | --- |
| | | Abs(log<sub>2</sub>FC) | | $\pi$ | | dN/dS (*M. huetii*) | | dN/dS (*R. communis*) | |
|  |  | B (CI) | p | B (CI) | p | B (CI) | p | B (CI) | p |
| (Intercept) |  | -1.64<br>(-1.65 – -1.62) | **<.001** | -1.92<br>(-1.92 – -1.92) | **<.001** | -1.60<br>(-1.61 – -1.58) | **<.001** | -1.40<br>(-1.41 – -1.39) | **<.001** |
| Au | PAR | 0.01<br>(-0.02 – 0.05) | .512 | -0.01<br>(-0.01 – 0.00) | .105 | 0.03<br>(-0.01 – 0.07) | .120 | 0.01<br>(-0.02 – 0.04) | .447 |
|  | SL | 0.05<br>(0.00 – 0.10) | **.037** | -0.01<br>(-0.02 – -0.00) | **.039** | 0.07<br>(0.01 – 0.12) | **.017** | -0.02<br>(-0.06 – 0.02) | .335 |
| SB | female | 1.37<br>(1.25 – 1.49) | **<.001** | 0.01<br>(-0.02 – 0.03) | .582 | 0.21<br>(0.06 – 0.36) | **.006** | 0.03<br>(-0.10 – 0.17) | .642 |
|  | male | 1.66<br>(1.57 – 1.75) | **<.001** | -0.02<br>(-0.04 – 0.00) | .066 | -0.04<br>(-0.14 – 0.06) | .423 | 0.08<br>(0.00 – 0.15) | **.046** |
| Interac<br>tions | PAR:<br>female | -0.07<br>(-0.33 – 0.20) | .633 | -0.04<br>(-0.10 – 0.02) | .234 | -0.38<br>(-0.64 – -0.11) | **.005** | -0.05<br>(-0.30 – 0.20) | .688 |
|  | SL:female | -0.31<br>(-0.61 – 0.00) | .052 | -0.02<br>(-0.08 – 0.05) | .612 | -0.13<br>(-0.48 – 0.22) | .463 | 0.16<br>(-0.11 – 0.43) | .237 |
|  | PAR:mal<br>e | 0.12<br>(-0.15 – 0.40) | .384 | 0.03<br>(-0.03 – 0.08) | .377 | -0.02<br>(-0.31 – 0.28) | .903 | -0.05<br>(-0.38 – 0.27) | .747 |
|  | SL:male | 0.03<br>(-0.27 – 0.33) | .824 | 0.09<br>(0.03 – 0.15) | **.004** | 0.13<br>(-0.32 – 0.58) | .562 | -0.06<br>(-0.32 – 0.20) | .645 |
| Observations |  | 10076 |  | 9345 |  | 4761 |  | 2993 |  |
| R2 / adj. R2 |  | .181 / .181 |  | .002 / .001 |  | .004 / .002 |  | .003 / .000 |  |
| AIC |  | 17367.804 |  | -13367.118 |  | 5774.396 |  | 914.510 |  |

**Table S10:** Linear models on Box-Cox-transformed measures of sex-biased gene expression for sex-biased genes of *M. annua* on their own (absolute log_2_FC difference between the sexes), nucleotide diversity (π), and pairwise d_N_/d_S_ (with reference to *M. huetii* and *Ricinus communis)*, as a function of sex linkage (SL: sex linked; PAR: pseudoautosomal; Au: autosomal), sex-bias (male: male-biased; female: female-biased) and their interactions.

| Predictors |  | Dependent Variables |  |  |  |  |  |  |  |
| --- | --- | --- | --- | --- | --- | --- | --- | --- | --- |
| | | Abs(log<sub>2</sub>FC) | | $\pi$ | | dN/dS (*M. huetii*) | | dN/dS (*R. communis*) | |
|  |  | B (CI) | p | B (CI) | p | B (CI) | p | B (CI) | p |
| (Intercept) |  | -0.44<br>(-0.56 – -0.32) | **<.001** | -3.25<br>(-3.35 – -3.15) | **<.001** | -1.15<br>(-1.25 – -1.05) | **<.001** | -1.52<br>(-1.68 – -1.37) | **<.001** |
| SB | male | 0.28<br>(0.13 – 0.44) | **<.001** | -0.09<br>(-0.21 – 0.04) | .171 | -0.20<br>(-0.32 – -0.08) | **<.001** | 0.06<br>(-0.12 – 0.23) | .531 |
| Au | PAR | 0.01<br>(-0.26 – 0.28) | .947 | -0.31<br>(-0.53 – -0.09) | **.007** | -0.21<br>(-0.37 – -0.04) | **.018** | -0.05<br>(-0.34 – 0.24) | .743 |
|  | SL | -0.33<br>(-0.64 – -0.01) | **.041** | -0.07<br>(-0.33 – 0.18) | .578 | -0.05<br>(-0.28 – 0.17) | .644 | 0.18<br>(-0.13 – 0.48) | .249 |
| Interac<br>tions | male:PAR | 0.15<br>(-0.24 – 0.54) | .443 | 0.23<br>(-0.07 – 0.53) | .126 | 0.25<br>(0.00 – 0.51) | **.049** | 0.00<br>(-0.48 – 0.48) | .995 |
|  | male:SL | 0.47<br>(0.03 – 0.90) | **.036** | 0.25<br>(-0.08 – 0.59) | .138 | 0.29<br>(-0.08 – 0.65) | .124 | -0.29<br>(-0.72 – 0.13) | .177 |
| Observations |  | 316 |  | 224 |  | 159 |  | 92 |  |
| R2 / adj. R2 |  | .098 / .083 |  | .049 / .027 |  | .088 / .059 |  | .028 / -.028 |  |
| AIC |  | 566.527 |  | 232.044 |  | 61.812 |  | 60.485 |  |

**Table S11:**
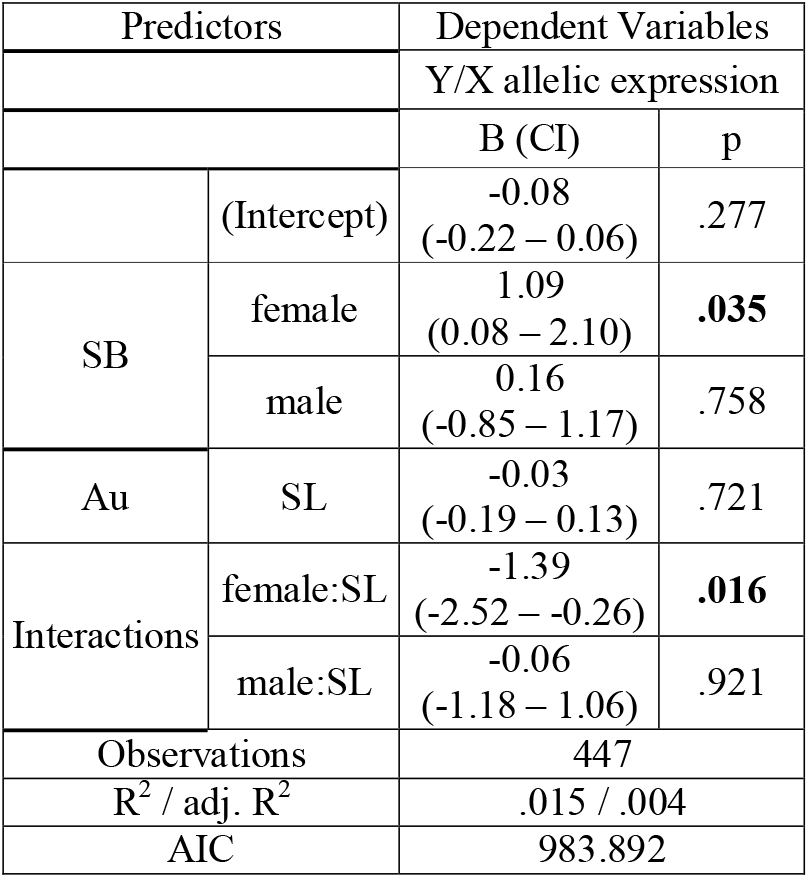
Linear models of Box-Cox-transformed measures of Y/X allelic expression and pairwise d_N_/d_S_ between X-and Y-linked alleles over all genes, as a function of sex linkage (SL: sex linked, PAR: pseudoautosomal, Au: autosomal), sex-bias (male: male-biased; female: female-biased) and their interactions.

**Table S12:**
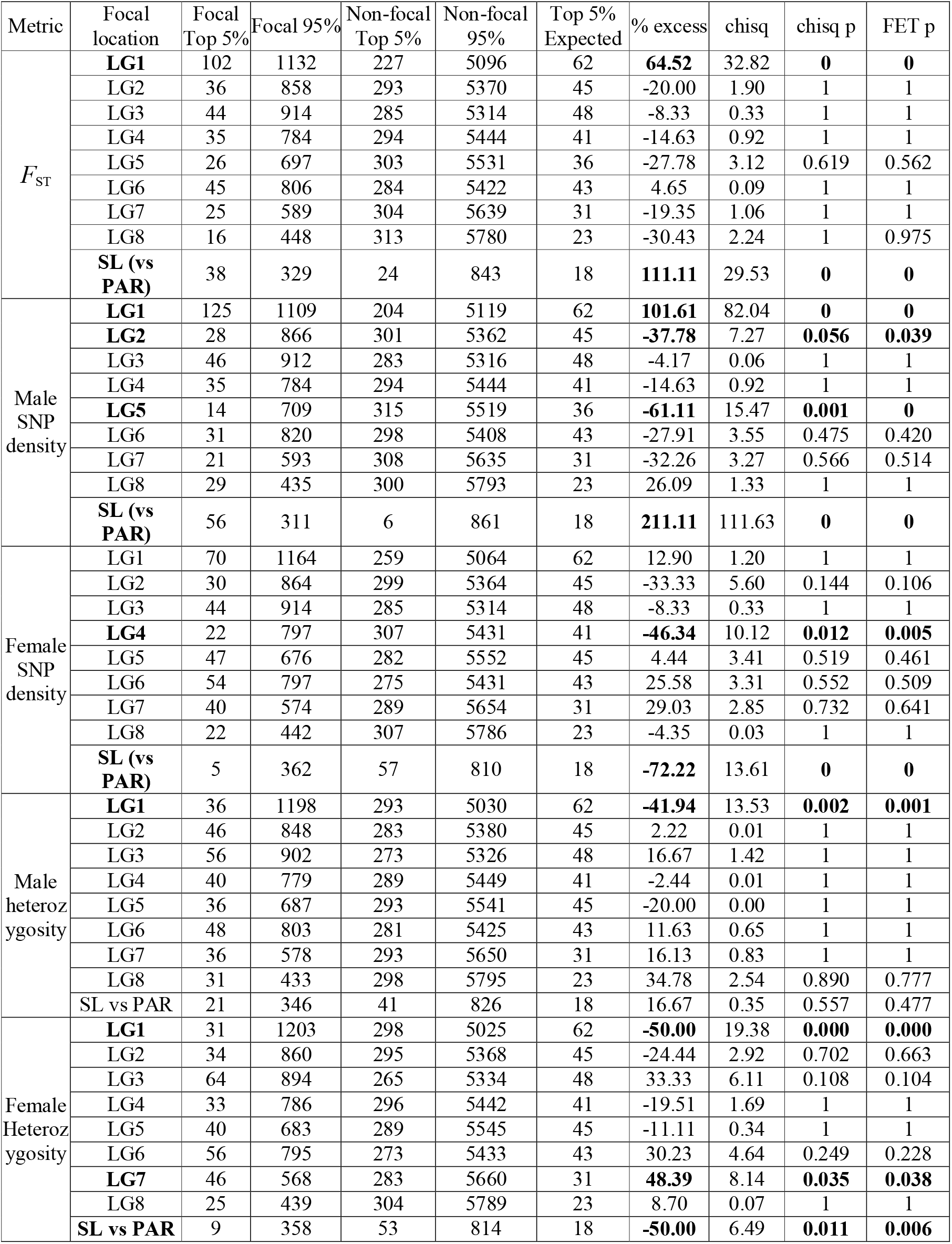
Summary of chi-squared and Fisher exact tests on the proportion of 5% outliers in *F*_ST_, male and female SNP density, and heterozygosity across the different linkage groups (focal being the LG shown in the second column), or within LG1 (focal being the sex-linked region). The p-values from comparisons between LGs have been adjusted for multiple testing and are indicated in bold when significant. SL = sex-linked.

| Metric | Focal location | Focal Top 5% | Focal 95% | Non-focal Top 5% | Non-focal 95% | Top 5% Expected | % excess | chisq | chisq p | FET p |
| --- | --- | --- | --- | --- | --- | --- | --- | --- | --- | --- |
| $F_{ST}$ | **LG1** | 102 | 1132 | 227 | 5096 | 62 | **64.52** | 32.82 | **0** | **0** |
|  | LG2 | 36 | 858 | 293 | 5370 | 45 | -20.00 | 1.90 | 1 | 1 |
|  | LG3 | 44 | 914 | 285 | 5314 | 48 | -8.33 | 0.33 | 1 | 1 |
|  | LG4 | 35 | 784 | 294 | 5444 | 41 | -14.63 | 0.92 | 1 | 1 |
|  | LG5 | 26 | 697 | 303 | 5531 | 36 | -27.78 | 3.12 | 0.619 | 0.562 |
|  | LG6 | 45 | 806 | 284 | 5422 | 43 | 4.65 | 0.09 | 1 | 1 |
|  | LG7 | 25 | 589 | 304 | 5639 | 31 | -19.35 | 1.06 | 1 | 1 |
|  | LG8 | 16 | 448 | 313 | 5780 | 23 | -30.43 | 2.24 | 1 | 0.975 |
|  | **SL (vs PAR)** | 38 | 329 | 24 | 843 | 18 | **111.11** | 29.53 | **0** | **0** |
| Male SNP density | **LG1** | 125 | 1109 | 204 | 5119 | 62 | **101.61** | 82.04 | **0** | **0** |
|  | **LG2** | 28 | 866 | 301 | 5362 | 45 | **-37.78** | 7.27 | **0.056** | **0.039** |
|  | LG3 | 46 | 912 | 283 | 5316 | 48 | -4.17 | 0.06 | 1 | 1 |
|  | LG4 | 35 | 784 | 294 | 5444 | 41 | -14.63 | 0.92 | 1 | 1 |
|  | **LG5** | 14 | 709 | 315 | 5519 | 36 | **-61.11** | 15.47 | **0.001** | **0** |
|  | LG6 | 31 | 820 | 298 | 5408 | 43 | -27.91 | 3.55 | 0.475 | 0.420 |
|  | LG7 | 21 | 593 | 308 | 5635 | 31 | -32.26 | 3.27 | 0.566 | 0.514 |
|  | LG8 | 29 | 435 | 300 | 5793 | 23 | 26.09 | 1.33 | 1 | 1 |
|  | **SL (vs PAR)** | 56 | 311 | 6 | 861 | 18 | **211.11** | 111.63 | **0** | **0** |
| Female SNP density | LG1 | 70 | 1164 | 259 | 5064 | 62 | 12.90 | 1.20 | 1 | 1 |
|  | LG2 | 30 | 864 | 299 | 5364 | 45 | -33.33 | 5.60 | 0.144 | 0.106 |
|  | LG3 | 44 | 914 | 285 | 5314 | 48 | -8.33 | 0.33 | 1 | 1 |
|  | **LG4** | 22 | 797 | 307 | 5431 | 41 | **-46.34** | 10.12 | **0.012** | **0.005** |
|  | LG5 | 47 | 676 | 282 | 5552 | 45 | 4.44 | 3.41 | 0.519 | 0.461 |
|  | LG6 | 54 | 797 | 275 | 5431 | 43 | 25.58 | 3.31 | 0.552 | 0.509 |
|  | LG7 | 40 | 574 | 289 | 5654 | 31 | 29.03 | 2.85 | 0.732 | 0.641 |
|  | LG8 | 22 | 442 | 307 | 5786 | 23 | -4.35 | 0.03 | 1 | 1 |
|  | **SL (vs PAR)** | 5 | 362 | 57 | 810 | 18 | **-72.22** | 13.61 | **0** | **0** |
| Male heteroz ygosity | **LG1** | 36 | 1198 | 293 | 5030 | 62 | **-41.94** | 13.53 | **0.002** | **0.001** |
|  | LG2 | 46 | 848 | 283 | 5380 | 45 | 2.22 | 0.01 | 1 | 1 |
|  | LG3 | 56 | 902 | 273 | 5326 | 48 | 16.67 | 1.42 | 1 | 1 |
|  | LG4 | 40 | 779 | 289 | 5449 | 41 | -2.44 | 0.01 | 1 | 1 |
|  | LG5 | 36 | 687 | 293 | 5541 | 45 | -20.00 | 0.00 | 1 | 1 |
|  | LG6 | 48 | 803 | 281 | 5425 | 43 | 11.63 | 0.65 | 1 | 1 |
|  | LG7 | 36 | 578 | 293 | 5650 | 31 | 16.13 | 0.83 | 1 | 1 |
|  | LG8 | 31 | 433 | 298 | 5795 | 23 | 34.78 | 2.54 | 0.890 | 0.777 |
|  | **SL vs PAR** | 21 | 346 | 41 | 826 | 18 | 16.67 | 0.35 | 0.557 | 0.477 |
| Female Heteroz ygosity | **LG1** | 31 | 1203 | 298 | 5025 | 62 | **-50.00** | 19.38 | **0.000** | **0.000** |
|  | LG2 | 34 | 860 | 295 | 5368 | 45 | -24.44 | 2.92 | 0.702 | 0.663 |
|  | LG3 | 64 | 894 | 265 | 5334 | 48 | 33.33 | 6.11 | 0.108 | 0.104 |
|  | LG4 | 33 | 786 | 296 | 5442 | 41 | -19.51 | 1.69 | 1 | 1 |
|  | LG5 | 40 | 683 | 289 | 5545 | 45 | -11.11 | 0.34 | 1 | 1 |
|  | LG6 | 56 | 795 | 273 | 5433 | 43 | 30.23 | 4.64 | 0.249 | 0.228 |
|  | **LG7** | 46 | 568 | 283 | 5660 | 31 | **48.39** | 8.14 | **0.035** | **0.038** |
|  | LG8 | 25 | 439 | 304 | 5789 | 23 | 8.70 | 0.07 | 1 | 1 |
|  | **SL vs PAR** | 9 | 358 | 53 | 814 | 18 | **-50.00** | 6.49 | **0.011** | **0.006** |

## Supplementary Information: Figures

**Figure S1:**
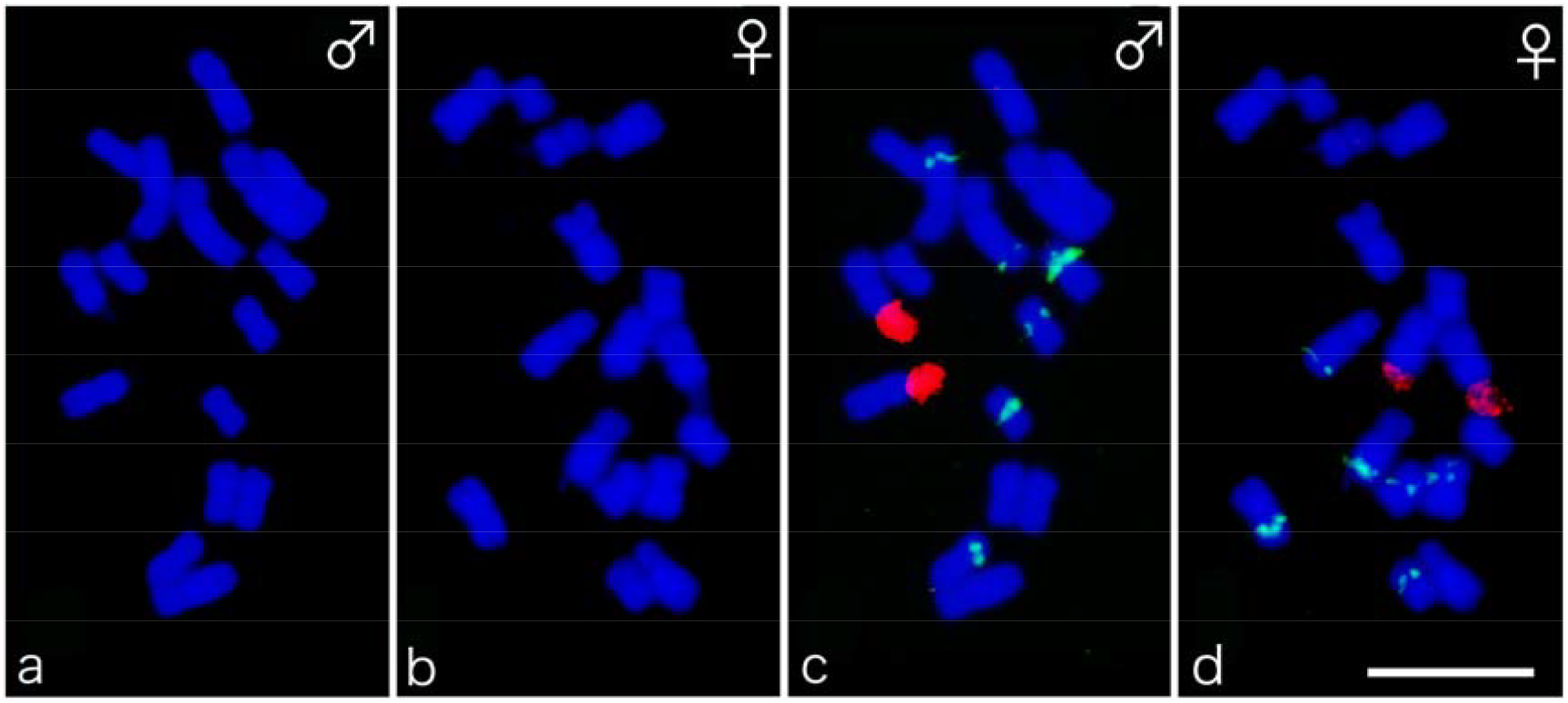
Male (a) and female (b) mitotic chromosomes of *Mercurialis annua* counterstained with DAPI (blue). Bicolor FISH of the same chromosome spreads is also shown (c and d). Hybridisation was performed with 45S rDNA (red) and 5S rDNA (green). Bar= 5 µm.

**Figure S2:**
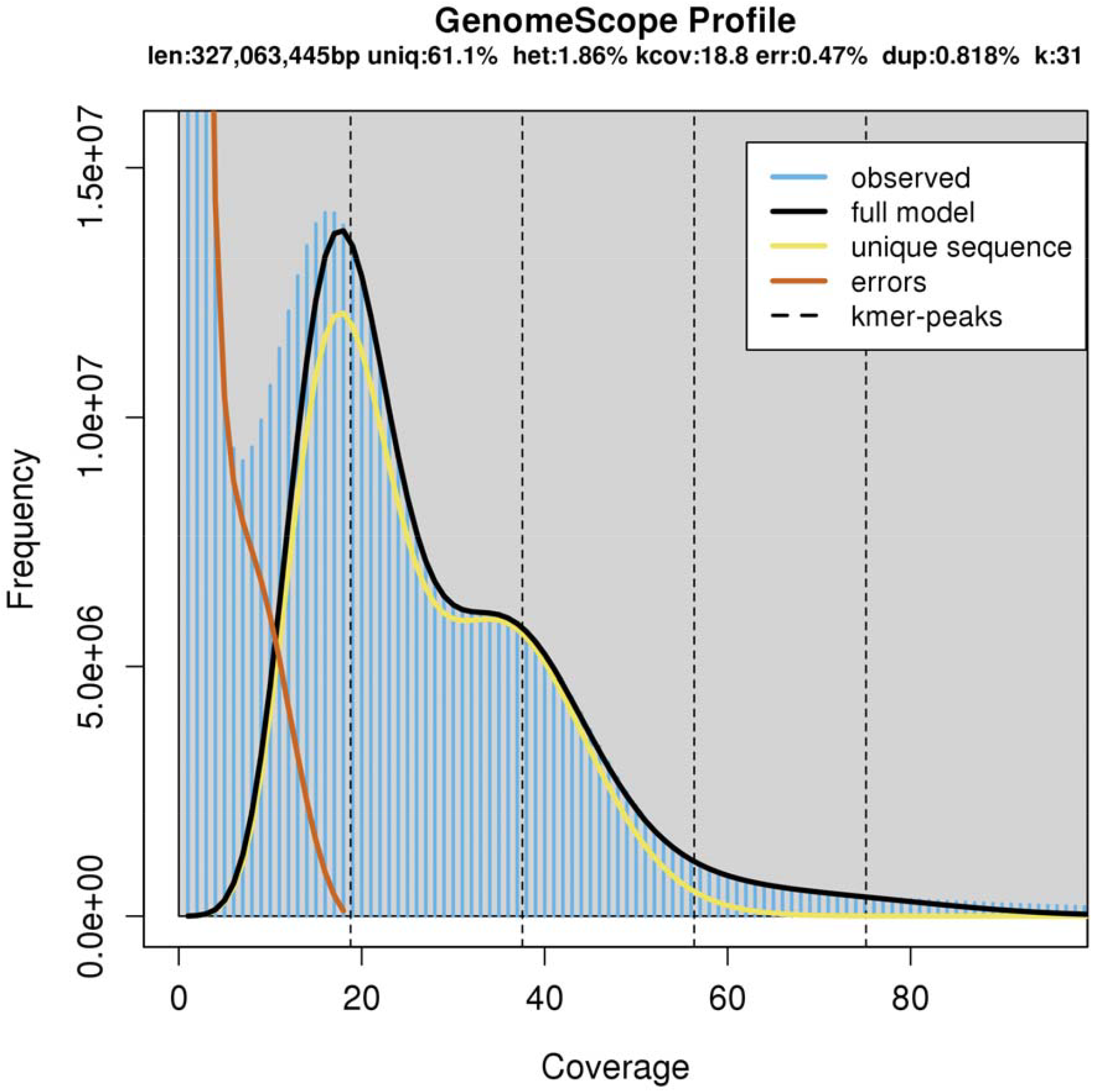
K-mer profile produced by GenomeScope.

**Figure S3:**
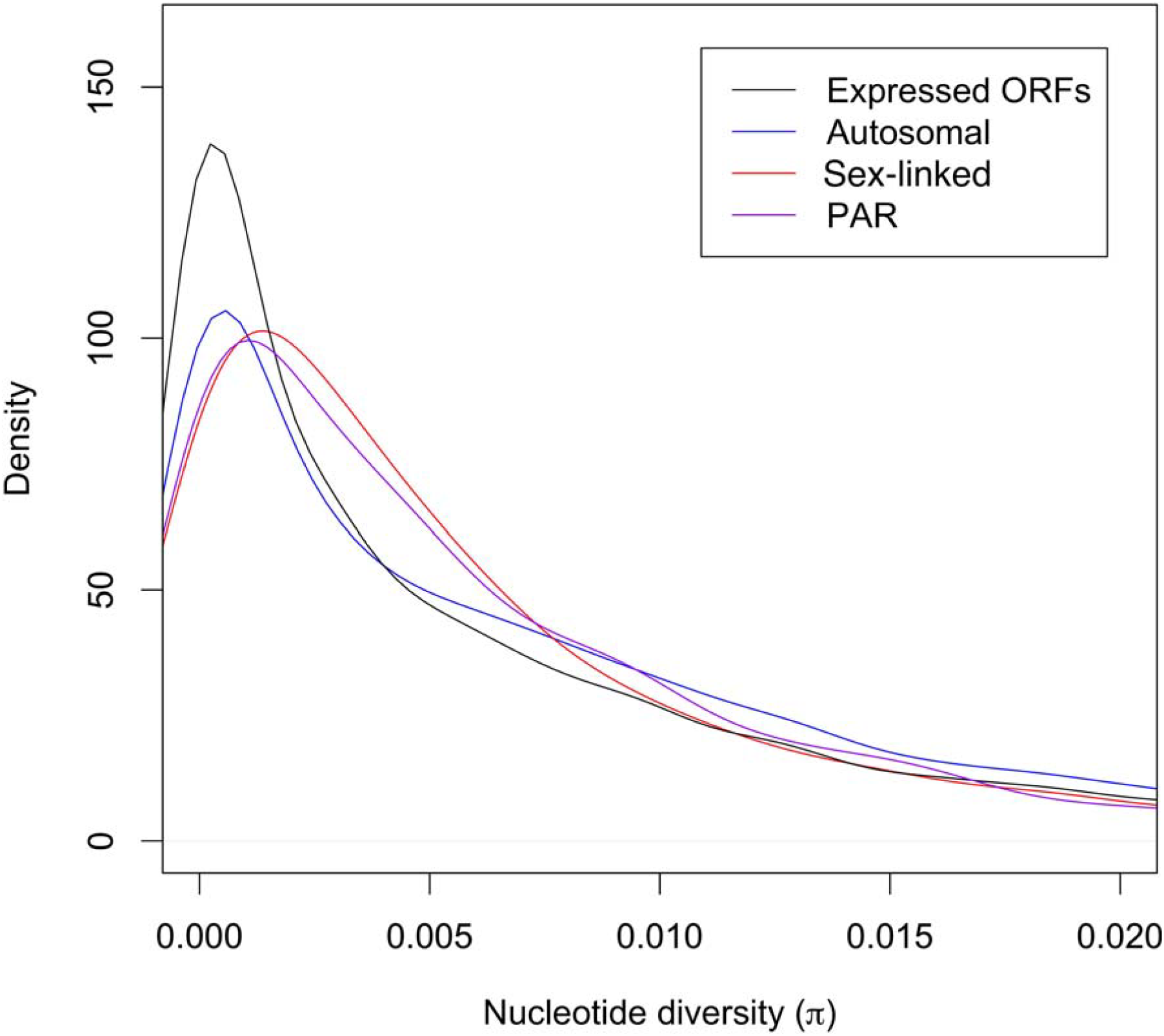
Density plot of nucleotide diversity (π) / kb across protein-coding genes for the X-linked, autosomal, PAR and expressed ORF bins. Density is calculated using a Gaussian kernel, such that the area under each curve sums to 1.

**Figure S4:**
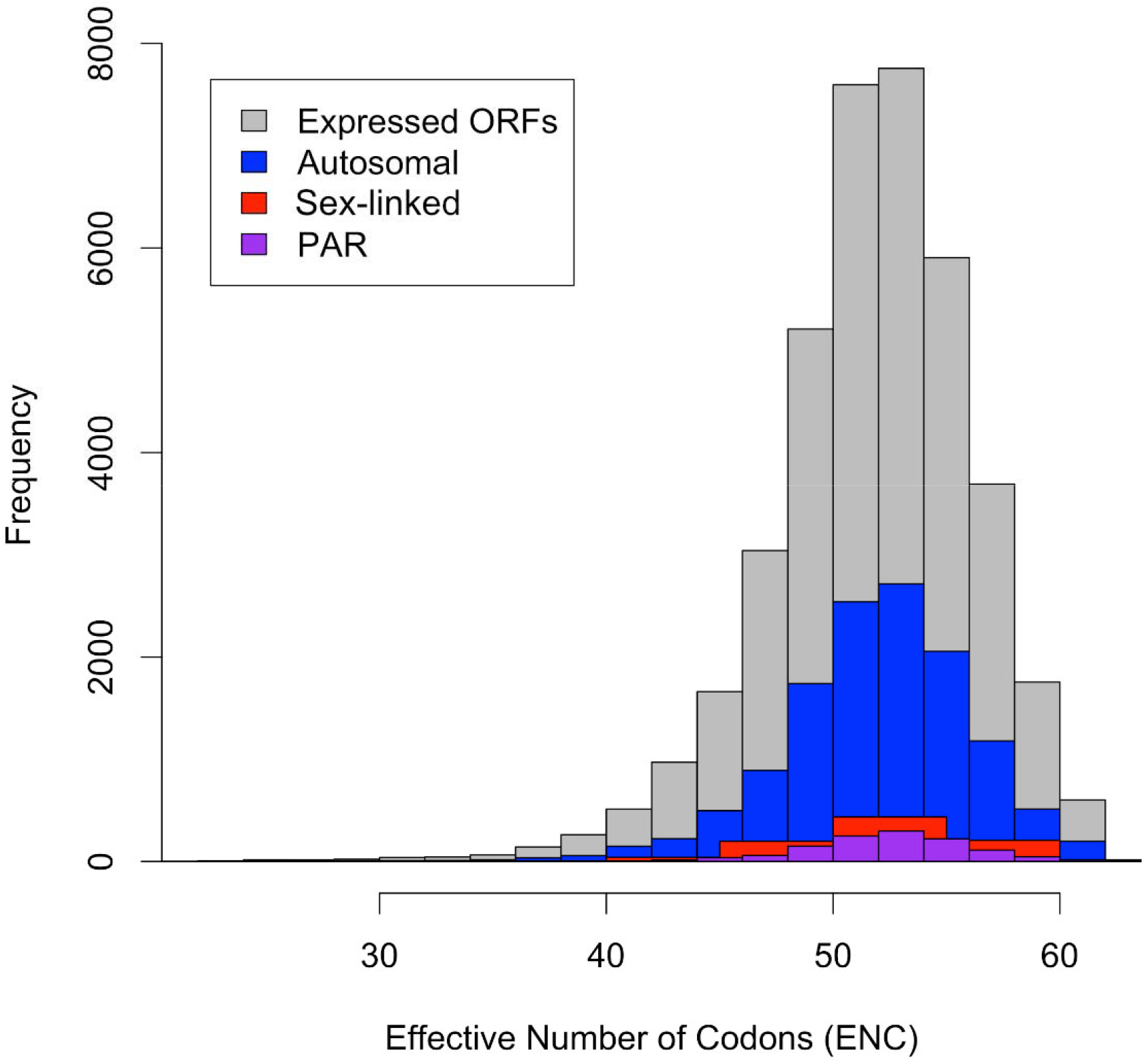
Histograms showing the effective number of codons per gene in the sex-linked, autosomal, PAR and expressed ORF bins.

**Figure S5:**
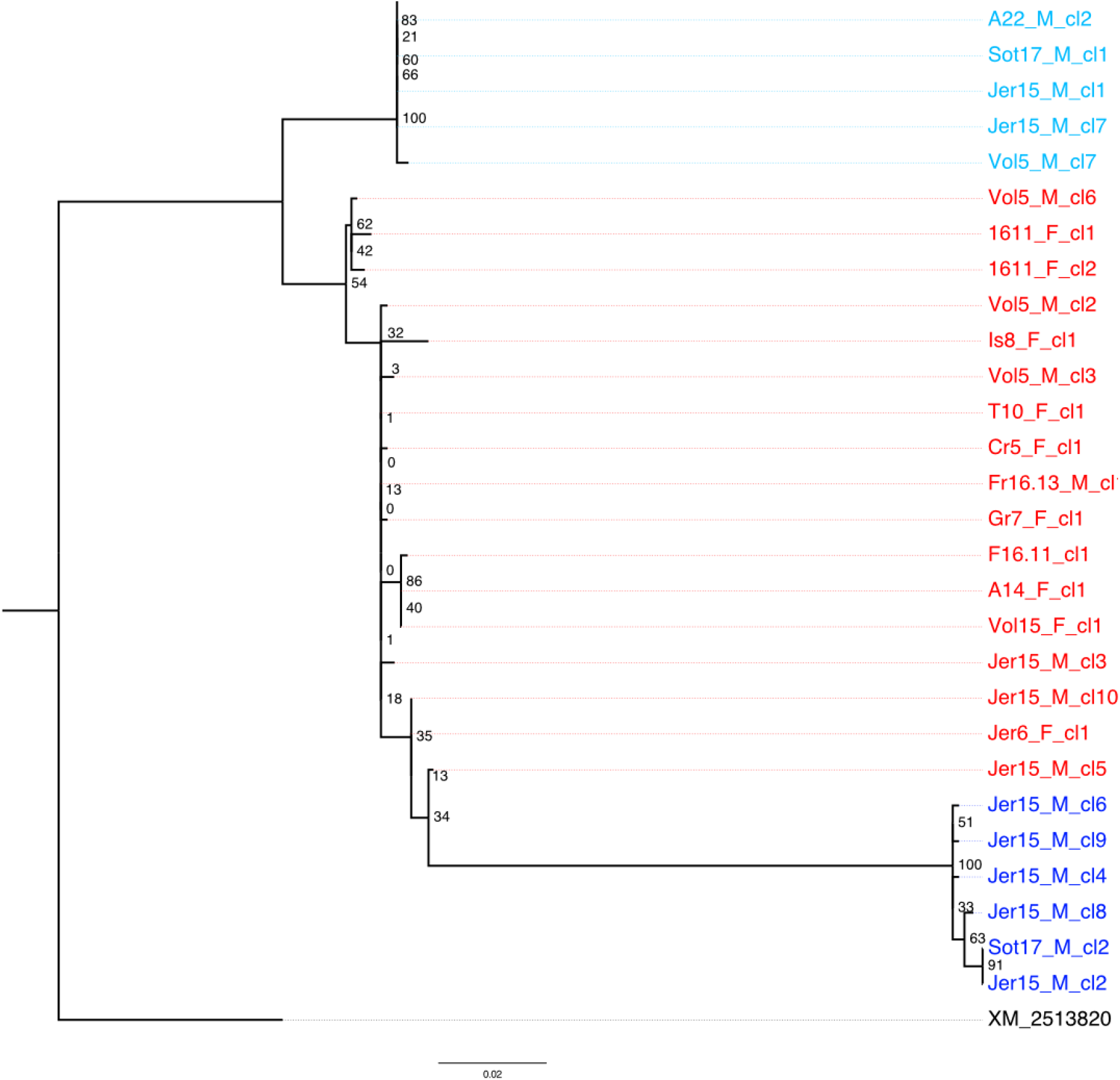
Phylogenetic tree of a sex-linked sequence inferred to have at least one premature stop codon in the Y-linked allele. Sequences marked in red amplify from both males and females, and are inferred to be on the X chromosome. The two groups marked in blue amplify from males only, sometimes from the same male indicating a duplication, and both have premature stop codons and other indels, which are more in the dark blue group. Numbers in black indicate node support from permutations. The outgroup is the *R. communis* homologue.

**Figure S6:**
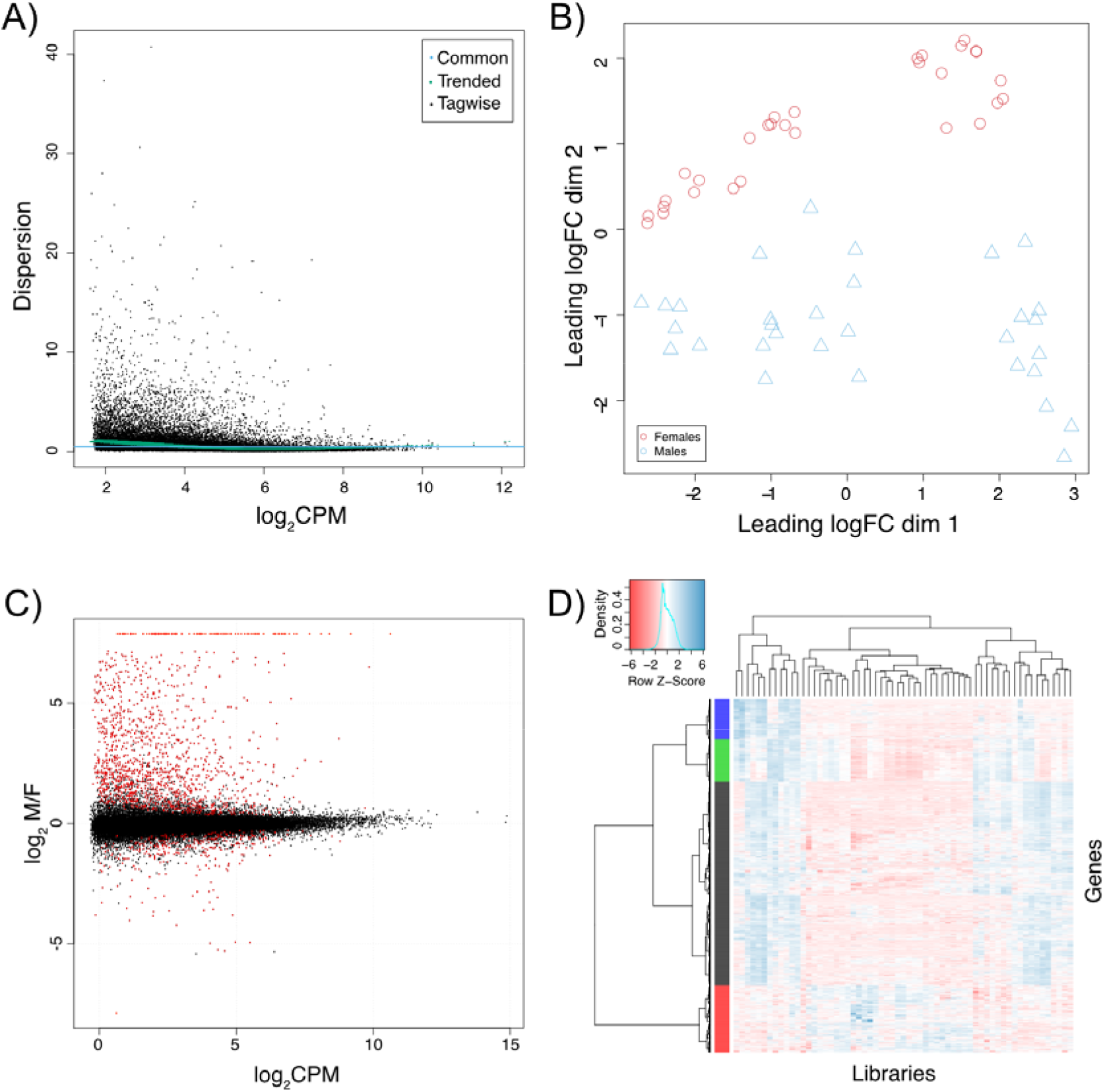
Summary of the differential expression analysis between males and females A) Dispersion plot, B) MDS plot with colour coding of male and female libraries, C) volcano plot indicating log fold change differences between males and females (differentially expressed genes are marked in red), D) heat map of the normalised counts in differentially expressed genes for each library.

## Supplementary Information: Files

**File S1**: Information on the mapped transcripts. Tab1: Information on expression, sex-bias, grouping in relation to recombination in males (SL, PAR, Au), position in male and female recombination map, linkage group assignment, pairwise d_N_, d_S_ compared to *M. huetii* and *R. communis* orthologs, first blastx hit name to swissprot, GO terms, relative expression of X and Y copy, nucleotide diversity. Tab2: Nucleotide diversity for pairs of transcripts between X/Y, *M. annua*/*M. huetii, M. annua*/*R. communis*. Tab3: Nucleotide diversity on transcripts with orthologs in both *M. huetii* and *R. communis,* used for estimating the Y branch length. Column names m1 and m2 refer to the model, lnL refers to their log likelihood. Topology 1 follows the species relationship, topology 2 is ((X, M. huetii)Y).

**File S2**: Fasta file of all sequences obtained from cloning and sequencing transcript 16827 from multiple individuals, and script used to generate the tree. The first letters in the sequence names are A: Antalya (TR), Jer: Jerusalem (IS), Vol: Volos (GR), 16: Montpellier (FR), T: Trabzon (TR), Sot: Southampton (UK). The following numbers identify the individual, and the sex is indicated by M (male) or F (female).

**File S3**: GO analysis summary for the sex-, male-and female-biased genes (one per tab). The first column abbreviates the main GO category -BP: Biological Process, CC: Cellular Component, MF: Molecular Function.

**File S4**: Full transcriptome annotation. The columns represent transcript name, best blastx hit to Swiss-Prot, and GO term annotation.

**File S5**: Transcriptome sequence. Includes the X and Y alleles, inferred from SEX-DETector, and the *M. huetii* transcriptome used.

**File S6**: Genomic contig sequence, and gff based on the transcriptome.

**File S7**: *M. annua* repeat library.

