## Supplementary material for "Early sex-chromosome evolution in the diploid dioecious plant *Mercurialis annua*": File S2 - tree stop codons: trees2.sh

```
#/bin/bash

#BSUB -L /bin/bash
#BSUB -o tree-output.txt
#BSUB -e tree-error.txt
#BSUB -J tree
#BSUB -u
#BSUB -N
#BSUB -n 4
#BSUB -R "span[ptile=4]"
#BSUB -R "rusage[mem=1000]"
#BSUB -M 1024288


module add SequenceAnalysis/MultipleSequenceAlignment/mafft/7.310;
module add Phylogeny/raxml/8.2.10;

mkdir /scratch/beegfs/monthly/pveltsos/raxml/${1}

cp ${1}.fasta /scratch/beegfs/monthly/pveltsos/raxml/${1}

cd /scratch/beegfs/monthly/pveltsos/raxml/${1}

mafft ${1}.fasta > ${1}_mafft.out

python ~/bin/fasta_to_phylip.py ${1}_mafft.out ${1}_mafft.phy

cp ${1}_mafft.phy ../

raxml -f d -p 12345 -N 100 -m GTRGAMMAI -s ${1}_mafft.phy -w /scratch/beegfs/monthly/pveltsos/raxml -T 4 -n ${1}

raxml -f d -p 12345 -N 1000 -m GTRGAMMAI -s ${1}_mafft.phy -w /scratch/beegfs/monthly/pveltsos/raxml -b 12345 -T 4 -n ${1}_b

raxml -f b -m GTRGAMMAI -z RAxML_bootstrap.${1}_b -t RAxML_bestTree.${1} -s ${1}_mafft.phy -w /scratch/beegfs/monthly/pveltsos/raxml -n ${1}_final -T 4

cp RAxML_bipartitions.${1}_final ${1}.tree

rm *${1}*RUN*

rm RAxML*${1}*

rm ${1}*phy*

rm ${1}_renamed.fa
```
